# Parallel alpine differentiation in *Arabidopsis arenosa*

**DOI:** 10.1101/2020.02.13.948158

**Authors:** Adam Knotek, Veronika Konečná, Guillaume Wos, Doubravka Požárová, Gabriela Šrámková, Magdalena Bohutínská, Vojtěch Zeisek, Karol Marhold, Filip Kolář

## Abstract

Parallel evolution provides powerful natural experiments for studying repeatability of evolution. Well-documented examples from plants are, however, still rare, as are inquiries of mechanisms driving convergence in some traits while divergence in others. *Arabidopsis arenosa*, a predominantly foothill species with scattered morphologically distinct alpine occurrences is a promising candidate. Yet, the hypothesis of parallelism remained untested. We sampled foothill and alpine populations in all regions known to harbour the alpine ecotype and used SNP genotyping to test for repeated alpine colonisation. Then, we combined field surveys and a common garden experiment to quantify phenotypic parallelism. Genetic clustering by region but not elevation and coalescent simulations demonstrated parallel origin of alpine ecotype in four mountain regions. Alpine populations exhibited parallelism in height and floral traits which persisted after two generations in cultivation. In contrast, leaf traits were distinctive only in certain region(s), reflecting a mixture of plasticity and genetically determined non-parallelism. We demonstrate varying degrees and causes of parallelism and non-parallelism across populations and traits within a plant species. Parallel divergence along a sharp elevation gradient makes *A. arenosa* a promising candidate for studying genomic basis of adaptation.

## Background

When and why does evolution result in a predictable outcome remain challenging questions in evolutionary biology. Parallel emergence of identical phenotypes in similar environments brings one of the most intuitive examples of the action of natural selection and provides independent biological replicates allowing for identifying the genetic basis and phenotypic outcomes of adaptation (1,2). In particular, parallel evolution of genetically-determined phenotypically distinct entities within a species, so-called ecotypes, provide valuable insights into processes of local adaptation and incipient ecological speciation in nature (3,4). In contrast to well-studied examples of ecotypic parallelism from animals such as fishes (5–8), snails (9,10) and stick insects (11,12), mechanisms driving parallel evolution in plants are largely unknown (but see refs. 13,14).

While directional selection driven by similar environment shall lead to convergence, alternative neutral and selective forces may counteract it, leading to larger genetic and phenotypic divergence among the derived ecotypes (reviewed in 3). Specifically, both stochastic processes such as genetic drift, varying intensity of gene flow and different pools of initial variation available for selection as well as deterministic processes such as selection in response to locally distinct micro-environments and genetic architecture of the traits can lead to non-parallel patterns in phenotypic variation (15,16). Consequently, there is no clear-cut border between parallel and non-parallel phenotypes, and the extent of parallelism may vary over traits or particular sets of compared populations (8). Finally, apparent similarity across populations can reflect short-term environmentally induced traits (phenotypic plasticity), however, the relative contribution of the heritable vs. plastic forces in the manifestation of phenotypic parallelism remains unknown (17).

To infer the mechanisms driving parallel phenotypic evolution as well as the extent of variation in parallel vs. non-parallel response, multiple independent environmental transitions within a species provide powerful naturally replicated systems. Indeed, independently evolved animal ecotypes documented that a combination of both stochastic and deterministic processes led to a varying extent of parallelism (3). In contrast, such detailed inquiries in plants are scarce due to a virtual lack of well-established systems that involve multiple (N > 2) replicates of environmental transitions within a species. The limited evidence from the so far investigated systems, *Heliosperma pusillum* (Caryophyllaceae; five pairs of alpine-foothill ecotypes, 14,18) and *Senecio lautus* (Asteraceae; seven pairs of dune-heathland ecotypes, 13,19), suggest considerable morphological divergence among the independently formed ecotypes despite similar environmental triggers, perhaps as a result of genetic drift (18) or gene flow (19). However, to infer mechanisms driving parallel evolution in plants in general and its genetic basis in particular, additional genetically well-tractable plant systems are needed.

Alpine populations of *Arabidopsis arenosa* represent a promising system for addressing drivers and consequences of parallel evolution in a genomic and molecular genetic context of the well-researched *Arabidopsis* genus. The species thrives mostly in low to mid-elevations (up to ∼ 1000 m a.s.l.) of Central and Eastern Europe, but scattered occurrences in treeless alpine habitats (∼ 1500– 2500 m a.s.l.) have been described from four mountain regions in the floristic literature (20–22). Alpine environments are generally well-poised for inquiry of plant parallel adaptation. On one hand, high elevations pose a spectrum of challenges to plant life, potentially triggering directional selection, such as freezing and fluctuating temperatures, strong winds, increased UV radiation and low partial pressure of carbon dioxide. Such pressures are believed to jointly trigger emergence of distinct alpine phenotypic ‘syndromes’ (e.g., contracted rosette plants, dense cushions, giant rosettes) that have been recurrently formed in distinct mountain ranges throughout the world (23–25). Indeed, alpine *A. arenosa* constitutively exhibits a distinct morphotype characterised by lower stature, less-lobed and thicker leaves, larger flowers and wider siliques (21). On the other hand, the island-like distribution of alpine habitats promotes parallel colonisation of individual alpine ‘islands’ by the spatially closest foothill populations (26). In line with this, a recent genomic study of *A. arenosa* (27) demonstrated geographical structuring of its range-wide genetic diversity, suggesting that the alpine ecotype might be of a polytopic origin. This hypothesis, as well as the extent of phenotypic parallelism and its plastic vs. genetic basis, however, remains untested.

In this study, we sampled multiple alpine and adjacent foothill *A. arenosa* populations covering all mountain regions known to harbour the alpine ecotype. Using genome-wide SNP genotyping we tested whether alpine environment within each mountain region has been colonised independently (regional clustering, ‘isolation-by-distance’ scenario) or if between-regional spread of the alpine populations took part (elevational clustering, ‘isolation-by-ecology’ scenario). Then, we combined analyses of field-sampled phenotypic data and a common garden experiment to uncover ecological and evolutionary mechanisms driving phenotypic parallelism. Specifically, we ask: (1) Had independent alpine colonisation triggered similar phenotypic transitions in distinct mountain regions? (2) Does the alpine phenotype remain divergent from the ‘typical’ foothill form in standardised conditions and, if so, which characters contribute to the genetically-determined alpine syndrome? (3) In contrast, are there traits exhibiting a rather opposite, non-parallel response?

## Methods

### Field sampling

*Arabidopsis arenosa* is a perennial outcrosser encompassing diploid and autotetraploid populations (27,28) that is widespread in foothill (colline to sub-montane) elevations across Central and Eastern Europe. Scattered occurrences of morphologically distinct populations have been recorded from four distinct mountain regions in Europe, sometimes treated as a separate species “*A. neglecta*”: Eastern Alps (20), Eastern (29), Southern (22) and Western Carpathians (21). While only tetraploid populations colonised the alpine stands in the former three regions, both diploid and tetraploid populations reached the alpine belt in the Western Carpathians. As the two cytotypes still occupy distinct alpine sub-regions there (diploids in Vysoké Tatry Mts. and tetraploids in Západné Tatry Mts.;, 30), we kept the ploidies as separate units for the sake of clarity. In sum, we hereafter refer to five regions: Niedere Taurern and surrounding foothills of the Eastern Alps (NT region, tetraploid), Rodna Mts. and adjacent regions of Eastern Carpathians (RD, tetraploid), Făgăraș Mts. in Southern Carpathians (FG, tetraploid), Vysoké Tatry Mts. and adjacent foothill diploid populations in Western Carpathians (VT, diploid), Západné Tatry Mts. and adjacent foothill tetraploid populations in Western Carpathians (ZT, tetraploid).

In each mountain region we sampled multiple populations from foothill habitats (semi-shaded rocky outcrops, screes and steep slopes; ‘foothill ecotype’) and from alpine sites (screes and rocky outcrops above the timberline; ‘alpine ecotype’). Both ecotypes are separated by a clear distribution gap spanning at least 500 m of elevation which also corresponds with the timberline. We avoided sampling plants in riverbeds immediately below the alpine populations that could represent recent colonizers germinated from washed seeds of the originally alpine plants. We sampled adult individuals from a total of 58 populations - 30 from foothill and 28 from alpine habitats. Within each population, we collected on average 17 individuals at the full-flowering stage: a small part of fresh tissue for ploidy determination, leaf tissue desiccated in silica gel for genotyping and vouchers for morphometrics. We selected largest rosette leaf, second stem leaf from the base and one random flower from the terminal inflorescence and fixed them onto paper with transparent tape for detailed measurements of organ sizes; the entire individual was then press-dried. For each population we also sampled the following local environmental parameters at a microsite with abundant occurrence of *A. arenosa*: (i) vegetation samples (phytosociological relevés, each covering an area of 3 × 3 m) recording percentage of the area covered by herb layer and listing all vascular plant species and (ii) mixed rhizosphere soil samples from five microsites within the vegetation sample.

Ploidy level of each sampled individual was determined using flow cytometry as described by Kolář *et al.* (28).

### Experimental cultivation

In addition to field sampling, we established a common garden experiment to test whether the plants of alpine origin keep their distinct appearance when cultivated in the foothill-like conditions. We used seeds of *A. arenosa* collected from 16 natural populations overlapping with those sampled for field phenotyping and genotyping (except for pop. AA254 that was used in the experiment as a replacement of spatially close, < 10 km, pop. AA145 which was included in the field dataset). The populations represent four regions (NT, VT, ZT and FG) and two distinct elevations (two foothill and two alpine populations per region; see Table S1 for locality details). To minimise maternal effect of the original localities, we firstly raised one generation in growth chambers under constant conditions. The plants used for phenotyping were raised in the Botanical Garden of the University of Innsbruck (Austria) situated in the Alpine valley, i.e. in conditions resembling the foothill habitat. Phenotypic traits were collected on all plants in the full-flowering stage in an identical way as for the field sampling. For details on cultivation conditions, see Methods S1.

### Inference of parallel origins

We genotyped 156 individuals from 46 populations (2-8 individuals per population, 3.5 on average) using the double-digest RADseq protocol of Arnold *et al.* (31). For an additional 44 individuals from 11 populations genome-wide SNP data were already available from a genome resequencing study (27). Raw reads processing and filtration generally followed our earlier study (31); for details see the Methods S1.

We inferred population grouping using several complementary approaches. Firstly, we ran K-means clustering, a non-parametric method with no assumption on ploidy, in adegenet v1.4-2 using 1000 random starts for each K between 1 and 20 and selected the partition with the lowest Bayesian Information Criterion (BIC) value (32). Secondly, we used model-based method FastStructure (33). We randomly sampled two alleles per tetraploid individual (using a custom script) - this approach has been demonstrated not to lead to biased clustering in autotetraploid samples in general (34) and *Arabidopsis* in particular (27). We ran the FastStructure with 10 replicates under K=5 (the same number as for K-means clustering, representing the number of the regions) and additionally only for tetraploid individuals under K=4. Finally, we displayed genetic distances among individuals using principal component analysis (PCA) as implemented in adegenet v.2.1.1. For clustering analyses, we used random thinning over 150 kb windows (length of our RAD-locus) to reduce the effect of linkage and removed singletons resulting in a dataset of 4,341 SNPs. PCA, AMOVA and genetic distances (see below) were calculated using the full set of 103,928 filtered SNPs.

Additionally, we tested for parallel origin of alpine populations using coalescent simulations in *fastsimcoal v.26* (35) – an approach that is also suitable for autotetraploids and mixed-ploidy systems (36). We constructed multi-dimensional allele frequency spectra from genome-wide SNPs from a subset of sufficiently sampled populations, one alpine and one foothill per region (∼ 176 – 417 k SNPs per population, see Table S2 and Methods S3 for details). We compared all regions occupied by tetraploid populations (NT, RD, FG, ZT) in a pairwise manner. Taking into account single origin of the widespread *A. arenosa* tetraploid cytotype (27,31), we had not compared all diploid-tetraploid pairs but only the spatially closest diploid and tetraploid populations from Western Carpathians (VT and ZT) for which a complex reticulation relationships was suggested previously (30). Briefly, for each pair of regions, we compared the fit of our data with two competing topologies: (i) sister position of alpine and foothill populations from the same region (i.e. parallel origin) and (ii) sister position of populations from distinct regions but belonging to the same ecotype (i.e. single origin of each ecotype). For each topology we either assumed or not assumed secondary contact (between-ecotype gene flow) within each region, what resulted in a total of four scenarios per regional pair (Fig 1C).

**Fig. 1.**
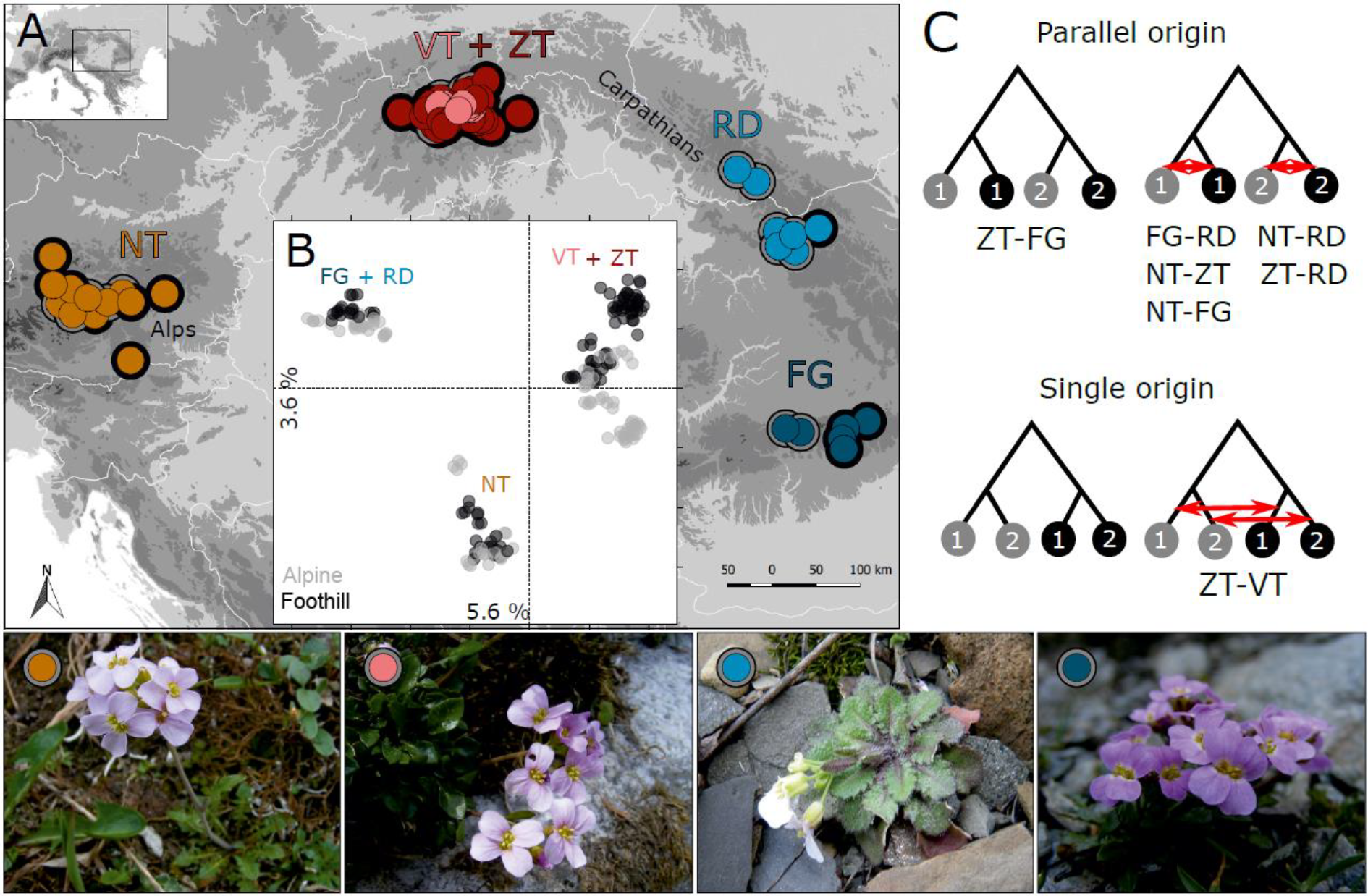
Parallel alpine differentiation of *Arabidopsis arenosa* in Central Europe inferred from genome-wide SNP variation. (A) Geographic distribution of sampled populations coloured by K-means clusters (circles with different colours) and classified according to the ecotype (black rim = foothill, grey rim = alpine). (B) Principal component analysis of the individuals coloured according to ecotype and labelled according to region (colour corresponding to pane A). (C) Four scenarios addressed by coalescent simulations; grey dots = alpine populations, black dots = foothill populations, numbers = regions, red arrows = migration events. The scenario with best fit to the observed allele frequency spectra for each pair of regions tested is indicated by its respective code below the scheme. Illustrative photos of convergent alpine *A. arenosa* ecotypes from four regions are presented at the bottom. Note that the VT region is occupied by diploids while all other regions are represented by autotetraploid populations.

We used hierarchical analysis of molecular variance (AMOVA) implemented in the R package pegas to test for genetic differentiation (i) among regions and among populations within regions and (ii) among ecotypes and among populations within each ecotype (both in the total dataset and in each region separately). We calculated nucleotide diversity (π) for each population with at least four individuals (N = 26) and pairwise differentiation (F_ST_) between these populations using custom python3 scripts (available at https://github.com/mbohutinska/ScanTools_ProtEvol). Finally, we tested for correlation between matrices of inter-population distances inferred (i) from SNP data (Nei’s distance calculated in StAMPP R package) and either (ii) from geographic location of populations (i.e., isolation-by-distance scenario) or (iii) from environmental parameters (Bray–Curtis distances based on eight environmental parameters, see below; isolation-by-ecology scenario) using Mantel tests with 9,999 permutations.

### Ecological data

To assess environmental differences among the populations, we combined broad-scale climatic parameters acquired from a database (SolarGIS, version 1.9, operated by GeoModel Solar, Bratislava, SK) with local conditions sampled at the original site of each population. First, we estimated the average values of three climatic variables: Precipitation, Temperature and Photosynthetic Active Radiation (PAR), over April, May and June, that correspond to the main growth period of *A. arenosa*. Second, we measured soil pH of each site by thermo-corrected electrode (WTW Multilab 540, Ionalyzer, pH/mV meter) at the Analytical Laboratory of the Institute of Botany (Průhonice, CZ). Finally, we inferred local environmental conditions from the vegetation samples: (i) total vegetation cover of herb layer and (ii) Ellenberg Indicators Values (EIVs) calculated from the species list using JUICE (37). EIVs provide estimates of environmental characteristics of the sites inferred from species composition based on expert knowledge (38). In total, we recorded 671 plant species of which 76 % have EIVs. We used only EIVs for Light, Nutrients and Moisture as the remaining EIVs (for Temperature, Continentality and Soil Reaction) overlapped with the above-described climatic and soil variables. In sum, our dataset of environmental parameters comprised the following eight variables: Precipitation, Temperature, PAR, soil_pH, Vegetation_cover, EIV_Light, EIV_Nutrients and EIV_Moisture. No pair of the parameters exhibited > 0.8 Pearson correlation (Table S3).

### Phenotypic traits

To test for parallel phenotypic response to alpine environment, we described the morphology of each plant sampled using 16 traits (999 / 223 vouchers in total scored for the field and common garden datasets, respectively). On each plant we measured 12 phenotypic characters describing the overall shape and size of both vegetative and reproductive organs (except for siliques which were not fully developed in the full-flowering stage which our sampling aimed at). To assess variation in shape of the organs independent of absolute size, we further derived four ratios (all characters are described and listed in Table S4). Missing values in the field dataset (1.5 % in total) were replaced by population means. For statistical analyses all characters except four (PL, PW, SL and SW), were log-transformed to approach normal distribution. No pair of traits was very strongly correlated (> 0.8, Table S3).

### Statistical analyses of ecological and morphological data

We used PCA calculated by base R function *prcomp* to visualise ecological differences among populations and morphological differences among individuals. Separate PCAs have been calculated on standardised (zero mean and unit variance) sets of (i) the eight climatic and local environmental variables and the 16 morphological traits recorded on the individuals collected (ii) in field and (iii) in a common garden experiment. Overall differentiation in environmental conditions and morphology of the ecotypes were tested by permutation multivariate analyses of variance (permanova). We first calculated Euclidean distances among population (environmental data) or individual (field and common garden morphological data) values using *dist* function and then ran a permanova test with *adonis2* function (number of permutations = 30,000) in R package vegan 2.5-4. In addition, we quantified the range of morphological variation of each ecotype by calculating disparity as the median distance between each individual and centroid of their corresponding ecotype in the ordination space using the R package dispRity.

To quantify morphological differentiation between pre-defined groups (ecotypes and regions) across all traits we ran classificatory discriminant analysis with cross-validation as implemented in Morphotools 1.1 (39). To assess relative contribution of individual morphological characters to the between foothill-alpine differentiation individuals we calculated a constrained ordination (linear discriminant analysis, LDA) in Morphotools. We calculated the discriminant analyses and permanova tests (i) for complete datasets and then (ii) separately for each region with ecotype as a factor to assess major drivers of foothill-alpine phenotypic differentiation within each region and (iii) separately for each ecotype with region as a factor to quantify variation in between-region phenotypic differentiation within each ecotype.

Finally, we ran generalised linear mixed models (GLM) for each of the 16 phenotypic traits to test for the effects of ecotype, region and their interaction (individual-based data with population as a random factor, *lme* function) in nlme package v3.1-137. Parallelism in foothill-alpine differentiation was considered for traits with significant effect of ecotype but non-significant ecotype × region interaction (lack of regionally-specific differences between ecotypes) that was revealed consistently in both field and common garden datasets. In turn, a trait with significant effect of ecotype × region interaction was indicative of non-parallelism (evidence for regional-specific differences). Finally, traits exhibiting significant ecotypic effect in the field dataset but not in common garden were considered as plastic with respect to alpine differentiation. To assess the degree of parallelism and non-parallelism in a more quantitative way, effect sizes of each factor and their interaction were estimated from a linear model using a partial eta-squared method (Etasq function) in R package heplots. Similarly as above, large ecotypic effect only was indicative of parallelism while strong contribution of the ecotype × region interaction implied non-parallel foothill-alpine differentiation in that particular trait (8). All the above analyses were run separately for the field and common garden datasets.

## Results

### Genetic structure and diversity

We confirmed the presence of tetraploid populations in four mountain regions (NT, ZT, RD, FG) while only diploids represented the Vysoké Tatry (VT) region in our sample. For the 200 RAD-sequenced individuals we gathered 103,928 SNPs of average depth 30× (1.37 % missing data). Non-hierarchical K-means clustering revealed that the populations cluster under K=5 (partition supported by the lowest Bayesian information criterion, Fig. S1) according to the geographic regions disregarding their alpine or foothill origin (i.e. the ecotype; Fig. 1). Populations from the spatially close VT and ZT regions, that differed by ploidy, were the single exception: here, the majority of the alpine diploid populations (from VT region) and some alpine tetraploid (ZT) populations formed a separate cluster distinct from the remaining Western Carpathian samples, regardless of ploidy. FastStructure run under K=5 supported this clustering and revealed clear separation of all groups but the VT and ZT groups, which were remarkably admixed (Fig. S2). Principal component analysis confirmed spatial, not ecotypic, clustering and revealed three main clusters: populations from the RD and FG regions were separated from the VT and ZT cluster along the first axis, while NT populations differentiated from the rest along the second axis (Fig. 1B).

Coalescent simulations demonstrated that parallel origin scenario is more likely than scenario assuming single origin of alpine ecotype and this result was consistent across all combinations of tetraploid populations regardless if we assumed subsequent gene flow between foothill and alpine populations within each region or not (median AIC difference between parallel and single origin scenarios ranged 2,550–115,250 and 3,829–252,144 over different pairs of regions, assuming and not assuming migration, respectively; Fig. 1C, Fig. S3). In contrast, for the spatially close diploid (VT) and tetraploid (ZT) populations the scenario involving sister position of alpine populations of both ploidies was preferred (median AIC difference from parallel scenario was 12,096 and 12,191, assuming and not assuming migration, respectively; Fig. 1C, Fig. S3). The difference in AIC between parallel origin scenarios assuming and not assuming gene flow was small (median AIC difference 28– 662) suggesting that allowing for subsequent gene flow between foothill and alpine populations within each region utmost slightly improved the fit of the model. In summary, genetic analysis coherently demonstrated polytopic origin of the alpine ecotype in four distinct mountain ranges (Alps - NT, Eastern Carpathians – RD, Southern Carpathians – FG and Western Carpathians – VT and ZT) and suggested reticulation between diploid (VT) and tetraploid (ZT) populations within the Western Carpathians. In order to account for the ploidy difference, we kept the VT and ZT populations separate in the following analyses. Additional analyses when the VT and ZT populations were merged into a single unit (Table S5), demonstrated that such alternative grouping does not lead to qualitatively different results with respect to the patterns in alpine-foothill differentiation.

Ecotypic differentiation explained a negligible proportion of genetic variance (3 %) while the five regions accounted for 20 % of variation (AMOVA analysis, see Table S6). Consequently, genetic differentiation by ecotype was non-significant within any of the mountain regions (Table S7, see also pairwise F_ST_ among populations, Table S8). Correspondingly, Mantel tests demonstrated high isolation-by-distance (rM = 0.39, p = 0.001) while isolation-by-ecology was non-significant (rM = 0.09, p = 0.129). The alpine colonisation was not accompanied by a reduction in genetic diversity, as populations belonging to both ecotypes exhibited similar nucleotide diversity overall (Wilcoxon test W = 77, p = 0.93) as well as within each mountain region separately (Table S7).

### Habitat differentiation

Environmental conditions of foothill and alpine sites significantly differed (permutational multivariate analysis of variance of populations, permanova, F = 16.92, p < 0.001; Fig. 2a,b). In contrast, populations from different mountain regions did not differ based on the environmental parameters recorded (F = 1.21, p = 0.30).

**Fig. 2.**
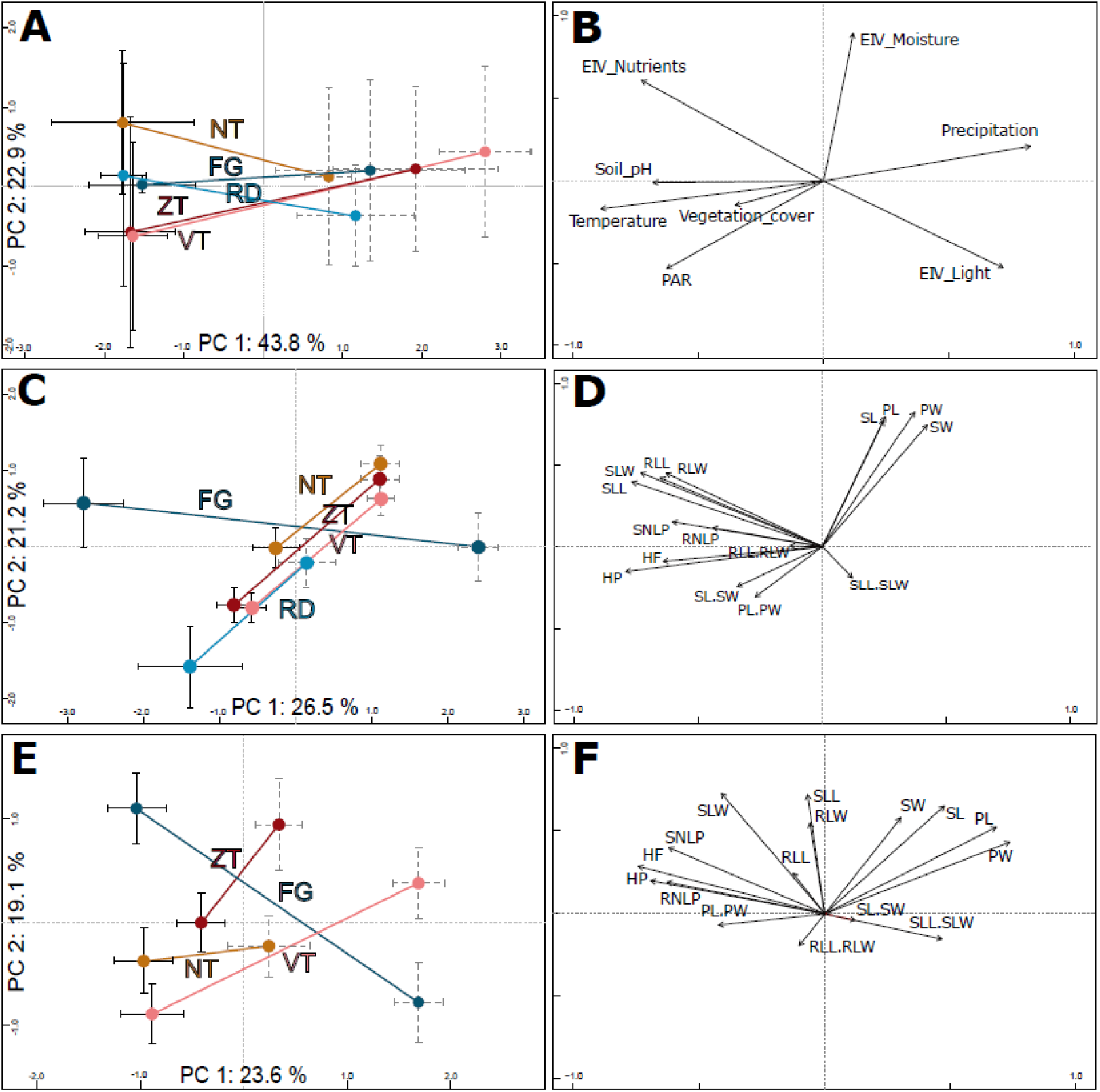
Parallel ecological (A, B) and field- (C, D) and common garden-based (E, F) morphological divergence of foothill and alpine populations of *A. arenosa* from the five mountain regions. The diagrams display results of principal ordination analyses (PCA) based on (A) eight environmental parameters of the original sampling sites of all populations with (C) 16 morphological traits collected in 58 wild populations, and (E) the same 16 traits sampled in a subset of 16 populations cultivated in a common garden. Centroids (dots) and standard error bars (black solid = foothill, grey dashed = alpine) are depicted for each region and ecotype; the ecotypes from the same region are connected by a line and coloured as in Fig. 1. The corresponding loadings of each environmental parameter (B) or morphological trait (D, F) are depicted. See Table S4 for explanation of individual codes of the traits.

### Morphological differentiation in natural populations

Field sampled alpine individuals were overall morphologically differentiated from their foothill counterparts as revealed by their separation in PCA (Fig. 2c, Fig. S4), high (87 %) classification success in the classificatory discriminant analysis and significant effect of ecotype in permanova (F = 49.25, p < 0.001). Morphological differentiation by ecotype was also significant for each region separately, with high classification success (89 %–100 % across regions, Table S7). Overall, morphological differentiation measured by disparity was significantly higher among the foothill than among the alpine individuals (F_1,997_ = 113.4, p < 0.001) suggesting that alpine individuals were morphologically more similar to each other than were their foothill counterparts.

Fourteen of the 16 scored characters significantly differentiated between the ecotypes, however, the level of parallelism remarkably differed across traits (Fig. 3). Stem height and traits reflecting flower size consistently varied between ecotypes across regions as demonstrated by significant effect of ecotype but non-significant ecotype × region interaction (GLM, Fig. 3, Table S9, Fig. S5) and strong contribution to the ordination constrained by ecotype (highest loadings to the discriminant axis in LDA Table S10). Overall, alpine plants, regardless of their region of origin, were shorter with larger calyces and petals (Table S4). In contrast, traits describing leaf size and shape were those with the strongest ecotype × region interaction indicating regionally-specific (i.e. non-parallel) morphological differentiation between ecotypes (Fig. 3). LDAs run separately for each region revealed such non-parallel response reflected distinct foothill-alpine differentiation in the FG region which was, apart from the height, most strongly driven by traits on leaves (LDA, Table S10). Such regionally-specific effect was primarily driven by the foothill FG morphotypes which is apparent from their very distinct position in the ordination space (Fig 2c) and higher distinctness of the foothill-FG than the alpine-FG populations when contrasted to populations from other regions belonging to the same ecotype (classification success as an FG group was 86 % vs. 77 % for the foothill vs. alpine populations, respectively).

**Fig. 3.**
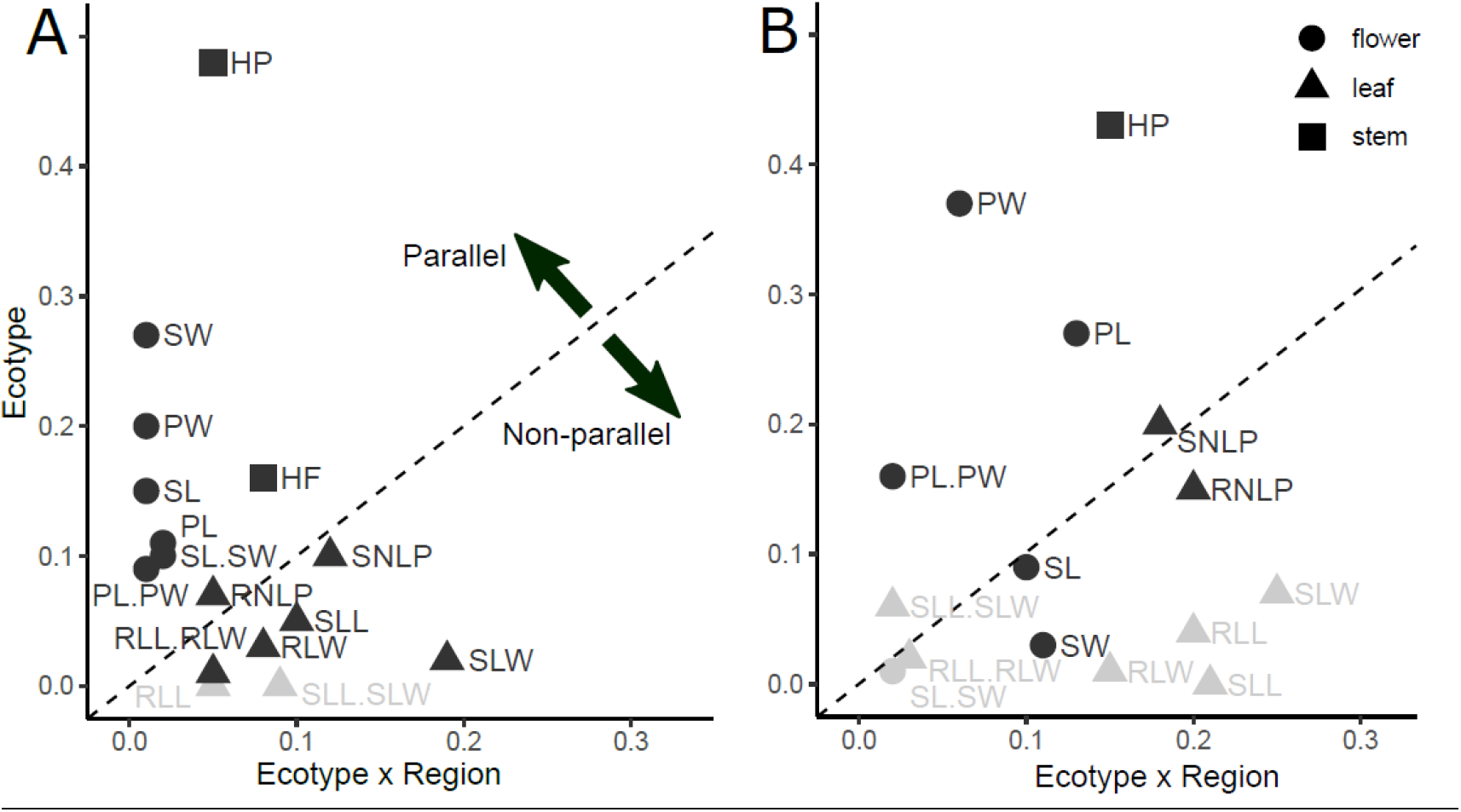
Variation in the extent of parallelism among individual phenotypic traits sampled in the field (A) and common garden (B). The extent of parallelism was estimated as effect sizes (Eta^2^) in linear model addressing the effect of Ecotype (foothill-alpine morphological divergence), Region (divergence between mountain regions) and their interaction, calculated separately for each trait across all populations sampled. The effect size of Ecotype (y-axis) shows the extent to which a given trait diverges predictably between ecotypes, i.e., in parallel, while the Ecotype × Region effect size (x-axis) quantifies the extent to which foothill-alpine phenotypic divergence varies across regions (i.e., deviates from parallel). Traits showing significant effect of Ecotype are marked in black. Points falling above the dashed line have a larger Ecotype effect (i.e. parallel) than Ecotype × Region interaction effect (i.e. non-parallel). The traits are categorized according to plant organ to which they belong (different symbols). See Table S4 for explanation of individual codes of the traits.

### Phenotypic variation in common garden

Overall phenotypic differentiation between plants originating from alpine and foothill environment remained highly significant even under two generations of cultivation in a common garden (a subset of 16 populations from four regions; Table S7, Fig. 2d). Differences between originally foothill and alpine populations were also significant within each region studied (permanova, Table S7), however, with markedly varying strength among the four regions examined, being weakest for NT plants (63 % classification success) and strongest for plants from FG region (95 %, Table S7). This implies that while alpine populations from some regions (FG, ZT in particular) keep their high morphological distinctness regardless of growth conditions, foothill and alpine populations from the NT region became more similar to each other when grown in a common garden. Morphological disparity of the originally foothill and alpine individuals was still significantly higher than that of their alpine counterparts (F_1,221_ = 30.96, p < 0.001).

Similarly to the field data, traits describing plant height and floral size contributed most strongly to the ecotypic differentiation across all regions (strong effect of ecotype but no ecotype × region interaction, Fig. 3; Table S9). On the other hand, ecotypic differentiation disappeared under common garden cultivation for nearly all leaf traits, with the exception of leaf lobe characters. However, leaf lobe traits also exhibited significant region × ecotype interaction due to their strong discriminative power between ecotypes in the FG region but not elsewhere (LDA, Table S10).

In summary, reduced stem height and larger floral organs showed the strongest parallelism in the genetic component of morphological differentiation across regions (significant effect of ecotype but non-significant ecotype × region interaction both in field and in the common garden; Fig. 3). On the other hand, leaf traits generally showed utmost regionally-specific discriminative power in the field (no effect of ecotype and/or non-significant ecotype × region interaction) and such regionally-specific discriminative effect mostly disappeared in a common garden, demonstrating plasticity in alpine-foothill differentiation for the majority of leaf traits. The only exception were leaf lobe traits that strongly discriminated foothill and alpine ecotype in one particular region (FG) constantly under both field and common garden conditions, demonstrating genetically determined non-parallelism.

## Discussion

Here, we combined a survey of genetic, ecological and morphological variation in natural populations with a common garden experiment to demonstrate multi-parallel ecotypic differentiation in a wild *Arabidopsis*.

### Parallel origin of the alpine ecotype

A mosaic distribution of ecotypes within a species’ phylogeny, with genetic diversity rather structured by geographical proximity (isolation-by-distance) than by environment (isolation-by-ecology), serves as evidence of repeated ecological divergence (40,41). The clustering, distance-based and coalescent analyses of genome-wide SNPs congruently demonstrated that the sampled *Arabidopsis arenosa* populations exhibit distinct regional clustering regardless of their alpine vs. foothill origin in four regions, i.e. supporting the ‘isolation-by-distance’ scenario. In line with biogeography (42,43), the major genetic groups corresponded to the four spatially well-defined and floristically distinct mountain regions: Eastern Alps (NT group) and Southern (FG), Eastern (RD) and Western Carpathians (VT+ZT groups). Genetic similarity of Western Carpathian diploids (VT) and tetraploids (ZT) has been detected previously and probably reflects recent origin of the tetraploid cytotype in the area and/or subsequent interploidy gene flow (27,30,31). However, due to distinct ploidy and spatial arrangement reducing the chance for extant gene flow among the VT-alpine and ZT-alpine populations, we considered the two regions as separate units in the following discussion for the sake of clarity.

Parallel origin of the alpine *A. arenosa* ecotype is consistent with previous range-wide studies of the species’ genetic structure (27,31). Taking into account spatial arrangement of populations and overall phylogeny of the species, colonisation of alpine stands from foothill populations likely underlies the observed foothill-alpine differentiation. Firstly, the foothill morphotype represents an ancestral state for the entire species; all three early-diverged diploid lineages of *A. arenosa* comprise only foothill populations (44). Secondly, foothill populations are spread over the entire area of Central and Eastern Europe wherever suitable rocky habitats are available (28) while the alpine populations are generally rare and confined to isolated spots within the five mountain regions investigated. Presence of such morphologically distinct populations in other areas is highly unlikely given the floristic knowledge of European mountains (21,22,45). Such foothill-to-alpine colonisation is also in line with overall expectations about the postglacial history of alpine flora in general (46) and of parallel alpine ecotypic differentiation within a species in particular (47–49). Interestingly, the exactly opposite alpine-to-foothill colonisation holds for the only other comparable example of multi-parallel differentiation in alpine plants, *Heliosperma* (14,18), making the comparisons among these systems more challenging.

### Parallel ecotypic differentiation

Independent colonisation of alpine stands followed by environmentally driven selection should lead to similar phenotypic changes across the alpine populations. Alternatively, drift, divergent selection to locally-specific conditions and/or limited variation in the source populations would lead to regionally specific patterns of foothill-alpine differentiation (3). Independently of the region of origin, alpine *A. arenosa* populations exhibited consistently shorter stems and larger flowers demonstrating parallelism in typical traits associated with ‘alpine syndrome’ that are considered adaptive in alpine environments (25,50). Elevation gradients belong to the most frequently studied environmental gradients in plant evolutionary ecology since the rise of this field (51,52), however, cases of phenotypic parallelism demonstrated by a combination of genetic and experimental data are still very rare in plant literature (53).

Alternatively, morphological similarities among the alpine populations may also reflect plastic response towards similar environmental conditions. To separate both effects, we grew plants of alpine and foothill origin from four regions in a common garden. Originally alpine plants kept their overall distinct appearance even in standard conditions, thus corresponding to the original definition of an ecotype *sensu* Turesson (54). Importantly, the traits exhibiting the strongest parallelism in foothill-alpine divergence in the field were also highly distinctive in the common garden (stem height and flower size), supporting the hypothesis of parallel differentiation over phenotypic plasticity. In contrast, the regionally-specific ecotypic differentiation detected in the field samples disappeared for the majority of such traits in cultivation, suggesting they rather manifested a plastic response to regionally-specific conditions. Such plastic response may still represent a way how to cope with the local nuances of the challenging alpine environments (e.g., 55), such a hypothesis would, however, require follow-up experimental tests.

### The degree and sources of non-parallelism

Even in systems exhibiting generally strong and genetically determined parallelism, both neutral and selective processes may cause significant non-parallel deviations in particular traits or populations (4,8). Although we observed significant non-parallelism within our set of traits, alpine *A. arenosa* populations showed overall phenotypic homogeneity that was relatively higher than that of their foothill ancestors (as indicated by relatively lower disparity of the alpine ecotype). This contrasts to remarkably divergent phenotypic outcomes of independent shifts along the gradient of elevation in the other multi-parallel plant system, *Heliosperma* (14). In this species, however, genetic drift in a small population of the derived (lower-elevation) ecotype was the likely driver of inter-population phenotypic divergence (18). In contrast, genetic drift likely had little impact on structuring diversity in alpine *A. arenosa*, as indicated by comparable genetic diversity across ecotypes and homogeneous population diversity in *A. arenosa* in general (27). Perhaps, gradual colonisation of alpine stands by populations pushed up by the continuously rising timberline during the Holocene, followed by survival in relatively large outcrossing and mostly polyploid populations, ensured the maintenance of large genetic diversity in the alpine ecotype.

For most of the traits exhibiting non-parallelism in the field data, ecotypic differentiation was no longer present under cultivation in the common garden, demonstrating phenotypic plasticity is the likely major driver of non-parallelism in our data. Only in one case (FG region) an additional trait (leaf lobes) discriminated foothill and alpine populations both in the field and in the common garden suggesting genetic determination of this component of non-parallelism. We consider a neutral scenario that initially divergent variation of the source (foothill) populations is likely responsible for this difference. Notably, the FG-foothill populations were more strongly divergent from the other foothill regions than were their FG-alpine counterparts from the other alpine populations. Emergence of non-parallel traits in response to local adaptation is less likely due to overall lower phenotypic variation among the alpine populations and very similar environmental conditions across alpine sites, speaking against strong selection triggered by locally-specific conditions. As parallelism informs about the role of selection, its detection is usually the prime aim of most studies (56), although the regionally-specific deviations provide valuable information on additional evolutionary processes affecting the predictability of evolution (8). Such inquiry, however, requires the study of multiple population pairs that evolved in parallel – a case that has been so far documented in only a handful of plant species (14,19).

### Conclusions

Our study demonstrated that spatially isolated alpine environments harbour populations exhibiting remarkable phenotypic parallelism that is manifested by traits that are associated with alpine adaptation in general. To which extent divergent selection shaped the differentiation and whether such phenotypic parallelism has also convergent genetic underpinnings, remains an open question, currently being followed-up by our ongoing study of genome-wide SNP variation in a subset of populations from each region. Indeed, alpine *A. arenosa* is a particularly well-suited model for following such questions due to negligible neutral divergence between ecotypes within each region and lack of strong bottleneck, both of which mitigate potentially confounding signals of past demographic events. This, complemented by the genomic and genetic tractability of the *Arabidopsis* genus in general and this species in particular (highly variable outcrosser) make *Arabidopsis arenosa* a highly promising model for studying the genomic basis of parallel adaptation in plants.

## Acknowledgements

The authors thank Eliška Záveská, Magdalena Lučanová, Jana Mořkovská, Jana Smatanová, Peter Schönswetter, Karl Hülber, Stanislav Španiel, Jakub Hojka, Daniel Bohutínský, Jindřich Chrtek, Klára Kabátová, Frederick Rooks and Martin Kolník for help with fieldwork. Erwann Arc, Dominik Kaplenig and Ilse Kranner kindly provided plants cultivated in common garden. Computational resources were provided by the CESNET LM2015042 and the CERIT Scientific Cloud LM2015085. The project was funded by Czech Science Foundation (project 19-06632S to KM, and 17-20357Y to FK), Charles University (GAUK project 708216 to AK) and Research Council of Norway (FRIPRO Mobility fellowship 262033 to FK). Additional support was provided by Ministry of Education Youth and Sports of the Czech Republic (7AMB18AT022 to GW).

