## Supplementary materials for "Parallel alpine differentiation in *Arabidopsis arenosa*"

**Methods S1*: Experimental cultivation conditions***

We firstly raised one generation of plants from each population in growth chambers under constant conditions. The field-collected seeds were germinated on a mixture of peat free potting compost and white river sand fraction 16-22 mm of neutral reaction in 2/3 ratio. Seedlings were re-potted at 2-4 leaves stage to 750 $cm^3$ pots with the same substrate, young rosettes were vernalized for 11 weeks (4 °C, 8/19 hrs day/night cycle) and then transferred to standard conditions in the chamber (21/18 °C, 16/8 hrs day/night cycle, light ~300 umol m-2*s-1). Each plant was hand-pollinated by a mixture of pollen from the same population (~ 14 flowering plants per population were used) for a period of one week. Ripe seeds (representing a genetically variable mixture of full- and half-sibs) were harvested and stored at dark and dry place at 4 °C.

The plants used for phenotyping were raised in the Botanical Garden of the University of Innsbruck, Austria (600 m a. s. l.; 47.26773 N, 11.37898 E). Seeds from min 10 maternal plants per population were sown in May 2018 on filter papers imbibed with 5 mM potassium nitrate and submitted to 4 days of cold stratification (4°C) before transfer to a growth chamber at 25 °C under constant light (~300 µmol $m^{-2}$*$s^{-1}$). Upon germination, seedlings were transferred to multi-pot trays using a mixture of peatless garden substrate and sieved sand of neutral reaction (1/1) and kept in an open-air greenhouse. Seedlings were then transplanted into individual pots (7 cm diameter, about 160 $cm^3$) filled with a mixture of rough-texture soil (5 parts of leaf compost, 2 parts of ground soil, 1 part of horticultural lava, 2 parts of coconut fibre, 2 parts of sand and 0.2 parts of stone powder) of neutral reaction (8/10), vermiculite (1/10) and sieved sand (1/10). All plants were kept in a sheltered outdoor place in the botanical garden over winter and the rosettes with set-up buds were brought back to the open-air greenhouse in mid-March 2019. Plants were watered regularly throughout the experiment. Finally, after four weeks the phenotypic traits were collected on all plants in the full-flowering stage.

**Methods S2: *Sequencing, raw data processing, variant calling and filtration***

***RADseq data***

One methylation-insensitive restriction enzyme (HpyCH4V) was used to generate blunt-end DNA fragments, followed by an A-tailing step combined with ligation of constructed adapters with T overhang and custom barcodes. Libraries were sequenced on an Illumina Hiseq 2500 platform using 125-bp paired-end reads at EMBL Genomics Core Facility (Heidelberg, Germany). Raw reads were demultiplexed, quality trimmed (> 30 Phred quality score) and mapped using BWA v. 0.7.3a on *Arabidopsis lyrata* reference v. 1.0.25 and the alignment was further processed with Picard Tools v. 1.119. The Genome Analysis Tool Kit v. 3.8-1 was used for simultaneous discovery and genotyping of SNPs and for calling of invariant sites following the recommended best practice (www.broadinstitute.org/gatk/). GATK performs SNP discovery and probabilistic genotype calling across all samples simultaneously, which is a more accurate than individual-based SNP calling. Namely, we used HaplotypeCaller module to call variants per individual with respect to its ploidy level (i.e., using the ploidy = 4 option which enables calling full tetraploid genotypes). Then we aggregated variants across all individuals by module GenotypeGVCFs. We retrieved SNP data for sites corresponding to the called RAD loci from a set of genome resequencing data of an additional 11 populations (mapped to the same reference and scored in the same way as described above, sequences available at Sequence Read Archive, SRA, code PRJNA484107). To do so, we extracted coordinates of the RAD loci, i.e. variant and invariant sites that were called in at least 30 % of individuals. Using these coordinates, we extracted the same sites from the filtered genomic vcf and combined both vcfs into a single file (CombineVariants). This merged raw vcf file containing the final set of 200 individuals from 57 populations was further filtered as follows: using GATK, we retained only biallelic sites that mapped to nuclear chromosome scaffolds with a minimum mapping quality of 40, which did not show mapping quality bias for the reads supporting the non-reference allele (keeping only variants with mapping quality rank sum test value higher than -12.5) and which were present in at least 80 % of our individuals at a sequencing depth of 8× or greater. In addition, we excluded potentially paralogous sites by masking genes that showed excess heterozygosity in a set of genome-resequenced populations from throughout the range of *A. arenosa*. We also masked sites that had excess read depth that we defined as 1.6× the second mode of the read depth distribution in the same genomic dataset.

**Methods S3:** ***Coalescent simulations***

We inferred the likely scenario of the origin of alpine populations using coalescent simulations performed in *fastsimcoal v.26*. We aimed to discriminate between single versus parallel origins of alpine ecotype, optionally assuming gene flow between ecotypes within each region, using population quartets involving one alpine and one foothill population from two regions. This strategy gave us four scenarios to test (single origin vs. parallel origin each with or without bi-directional between-ecotype admixture, see Fig. 1C), each of which was compared with observed data gathered for all six possible pairs of regions occupied by tetraploid populations, i.e. six quartets of populations from NT-RD, NT-FG, NT-ZT, RD-FG, RD-ZT, FG-ZT regions (i.e., 24 sets of simulations in total) and a pair of spatially closest diploids and tetraploids (VT-ZT). In order to keep the number of simulations realistic, the quartets involving diploids (VT) and other tetraploid regions were not simulated, as single origin of the tetraploid cytotype inferred previously makes origin of alpine tetraploids via independent polyploidisation from alpine VT diploids very unlikely.

Each model was fit to a multi-dimensional allele frequency spectrum calculated from putatively neutral four-fold degenerate SNPs. To gather robust allele frequency spectra, we acquired genome-wide SNP variation from a subset of sufficiently sampled populations (genome resequencing data, 8 individuals per population, 176,000 – 417,000 SNPs per population; Table S2), one alpine and one foothill per each region. We used populations for which genome resequencing data were available from a previous range-wide study (Monnahan et al. 2019, PRJNA484107) and complemented them with newly sequenced data (SRA project SUB6592572; see Table S2 for details). We extracted the four-fold degenerated SNPs (inferred based on *A. lyrata* annotation) using custom python scripts available at https://github.com/mbohutinska/ScanTools_ProtEvol, the allele frequency spectra, stored in **.dsfs* files, are attached as Supplementary data file S1. Read trimming, mapping, variant calling and filtration was performed as described above for RADseq data.

For each scenario and population quartet, we performed 50 independent fastsimcoal runs to overcome local maxima in the likelihood surface. We used wide range of initial parameters (effective population size, divergence times, migration rates; see the example **.est* and **.tpl* files provided in the Supplementary data file S1) and assumed mutation rate of 4.3e-8 inferred for *A. arenosa* previously (Arnold et al., 2015). We then extracted the best likelihood partition for each fastsimcoal run, calculated Akaike information criterion (AIC) and summarized them across the 50 different runs. The scenario with consistently lowest AIC values within particular population quartet was preferred (Fig. S3).

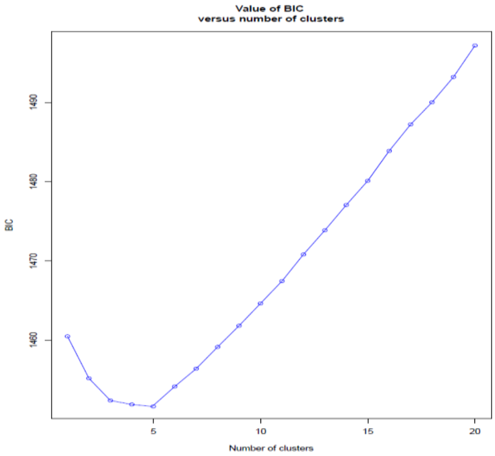

**Fig. S1** Selection of plausible number of clusters (K) in K-means clustering of SNP data. Bayesian information criterion (BIC) of K-means groupings from K=1 to K=20 summarized over 1000 random starts.

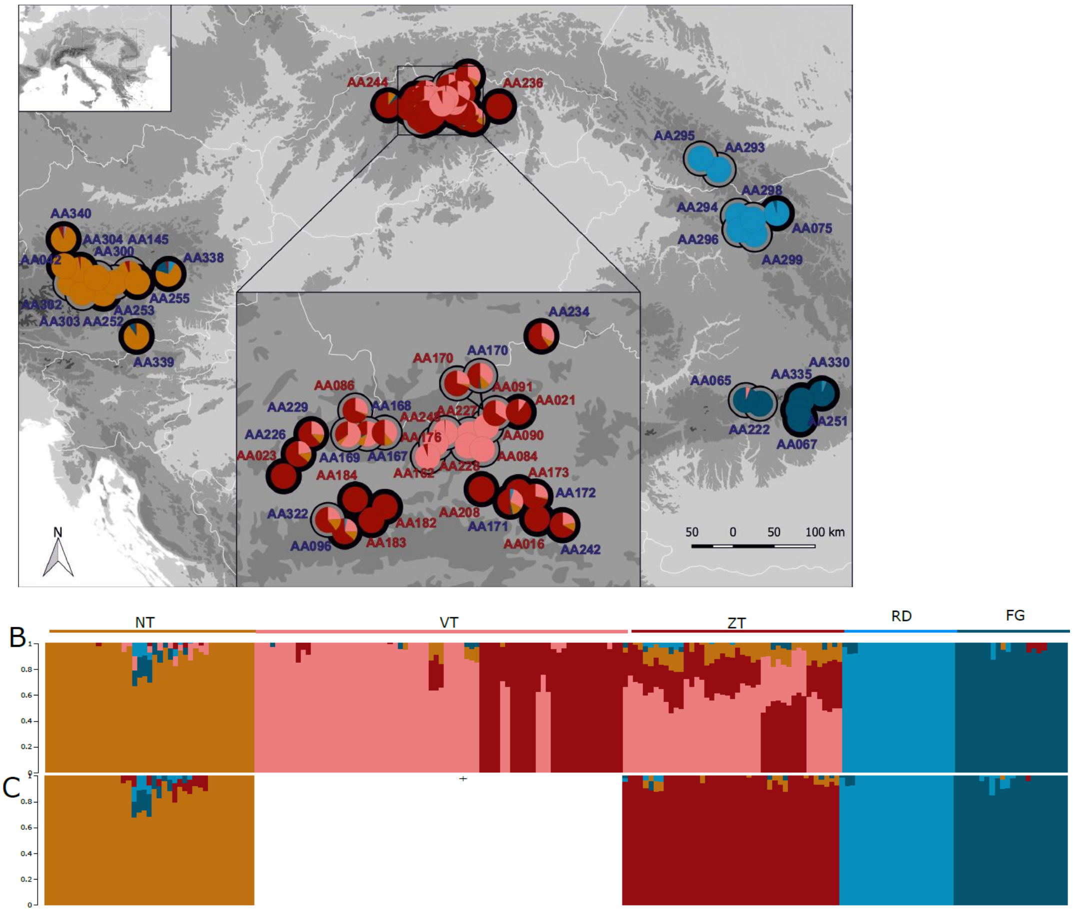

**Fig. S2** Genetic structure of alpine and foothill *Arabidopsis arenosa* identified by FastStructure. (A) Geographic distribution of the five genetic groups identified by FastStructure (different colours) and the two altitudinal ecotypes (different shading of circle margins: black = foothill, gray = alpine). Pie charts reflect assignment to different FastStructure groups, population labels are coloured by ploidy level (red = diploid, blue = tetraploid). The inset depicts detailed distribution in the Western Carpathians (a diploid VT and tetraploid ZT groups) (B) Posterior probability of assignment of all sampled *A. arenosa* individuals to the five groups as inferred by FastStructure under K=5. (C) Posterior probability of assignment of the tetraploid *A. arenosa* individuals to the four groups (four tetraploid regions only) as inferred by FastStructure under K=4.

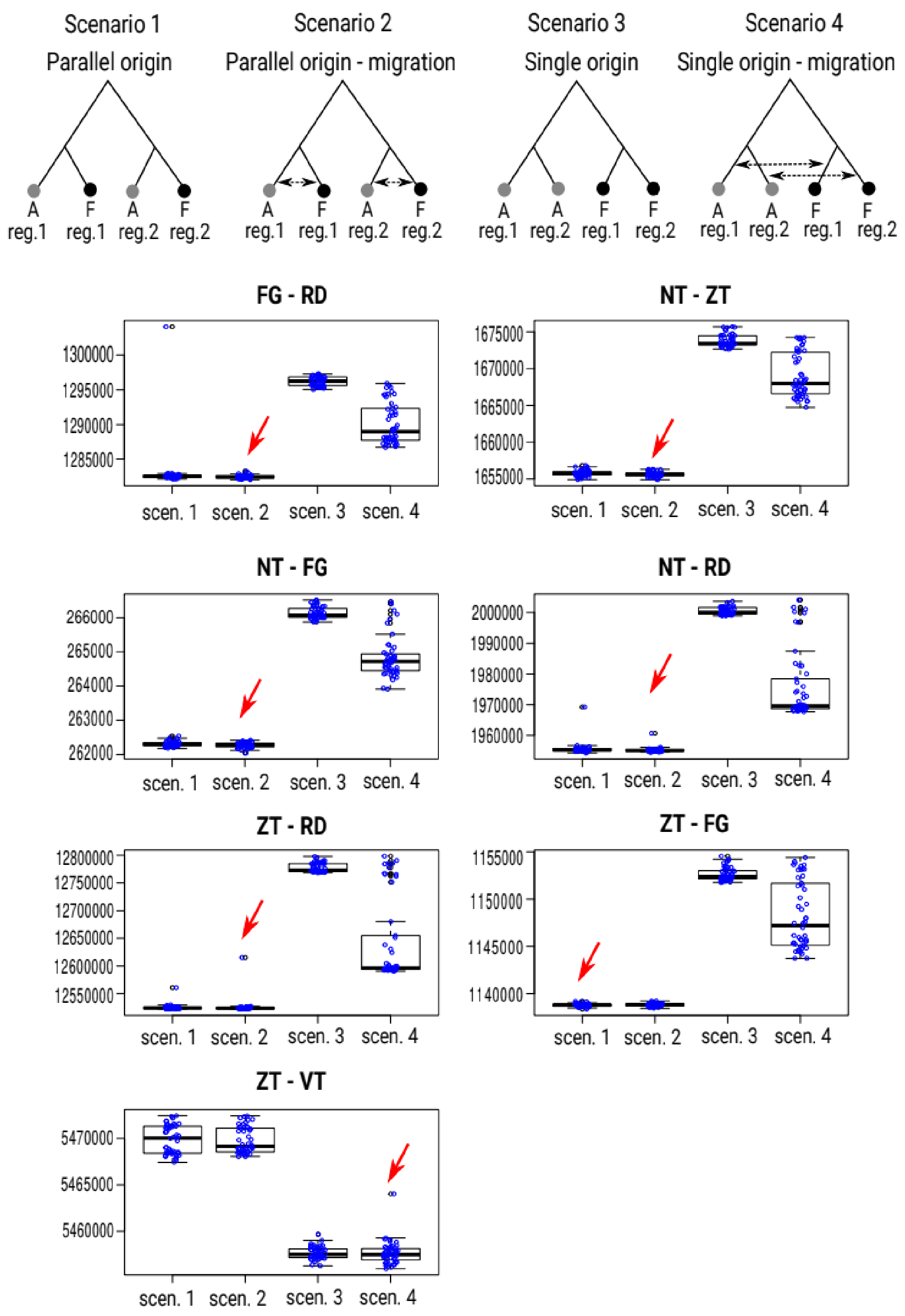

**Fig. S3** Comparison of Akaike information criteria (AIC) across four scenarios approximating the origin of the alpine ecotype of *A. arenosa* in pairs of mountain regions. We iterated alpine (A) and foothill (F) pops. from the following regions (reg.): FG, NT, RD, VT and ZT. Each scenario was simulated by 50 fastsimcoal2 independent runs, the corresponding distribution of the AIC values over these 50 runs (blue dots) is summarized by the boxplots. Red arrow highlights the most likely scenario with the lowest median AIC values. The topologies of the evaluated scenarios are depicted above the corresponding plots, black dot = foothill pop. (F), grey dot = alpine pop. (A).

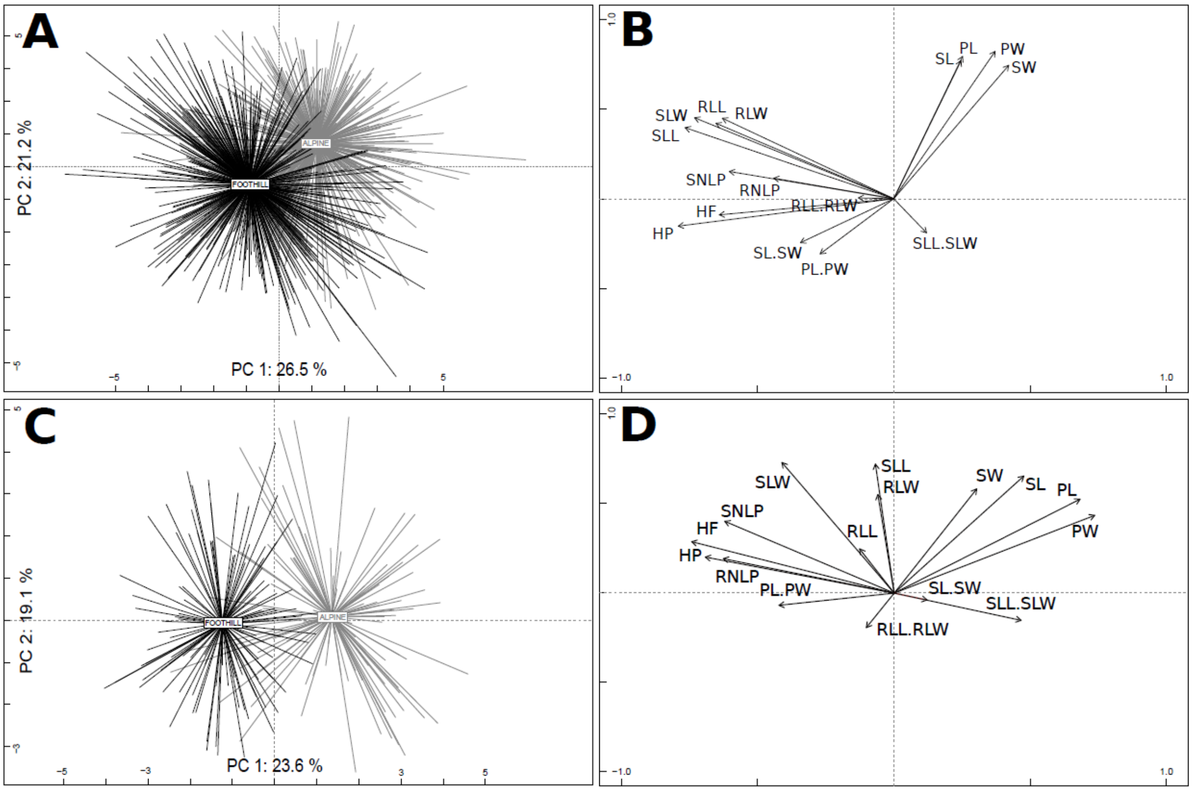

**Fig. S4** Morphological differentiation among foothill (black) and alpine (gray) individuals. Principal component analysis was run using 16 morphological traits based on (A, B) 999 individuals collected in field in five regions; (C, D) 223 individuals from the common garden experiment originating from four regions. Contribution of morphological characters for (B) plants collected in the field (D) plants from common garden experiment.

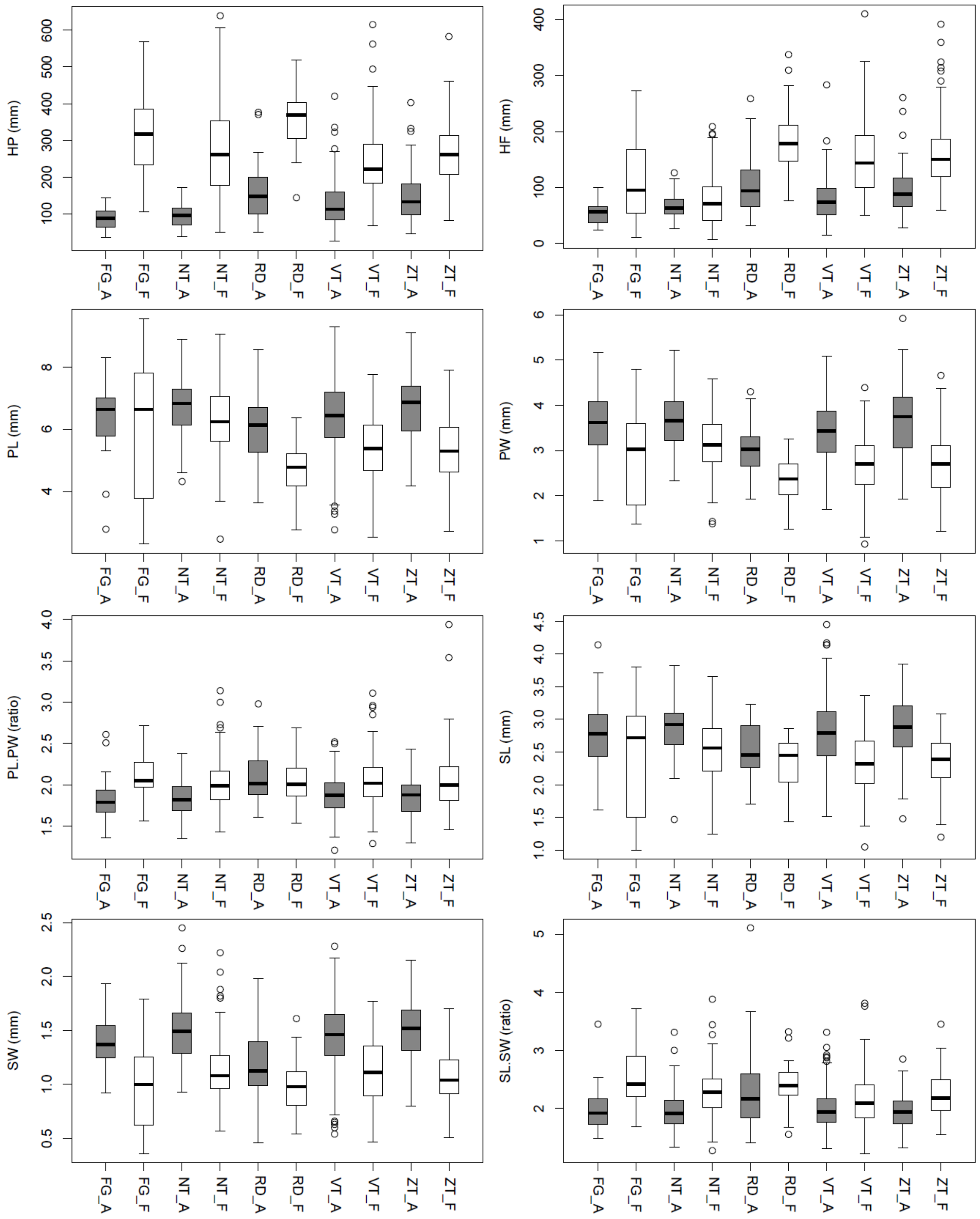
HP (mm)
HF (mm)
PL (mm)
PW (mm)
PL.PW (ratio)
SL (mm)
SW (mm)
SL.SW (ratio)
FG_A
FG_F
NT_A
NT_F
RD_A
RD_F
VT_A
VT_F
ZT_A
ZT_F

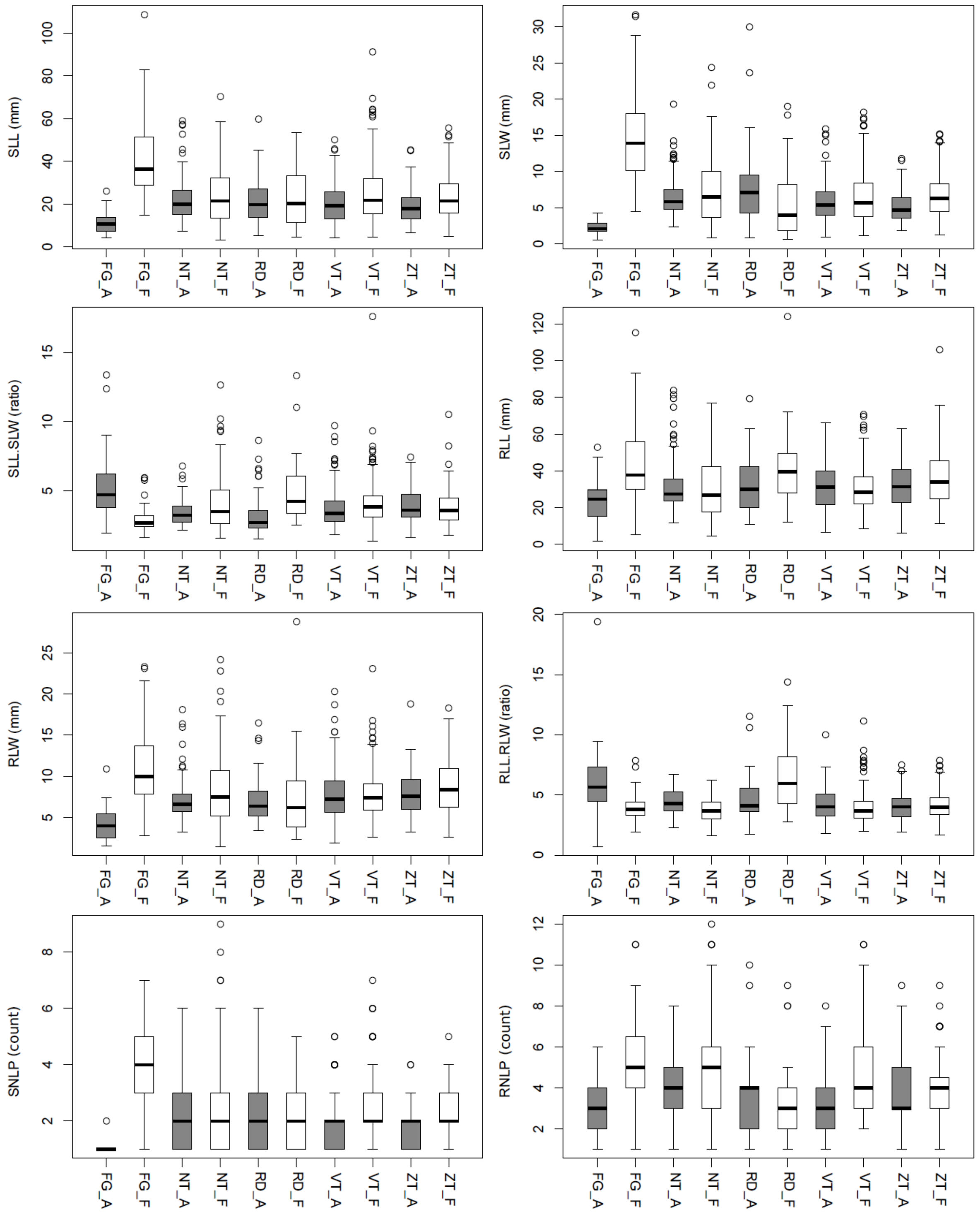

**Fig. S5** Distribution of trait values in foothill (F, white) and alpine (A, gray) groups of *A. arenosa* plants sampled in field in the five regions (FG, NT, RD, VT, ZT). Original values are provided in a Supplementary data file S3.

**Table S1** Details on ploidy, sampling location, habitat and genetic diversity of all populations sampled for the study.

| Population code | Ecotype | Region | Ploidy | Kmeans_group | N_morphology (field) | N genetic | Common Garden experiment | State | Locality | Collected | Latitude | Longitude | Elevation | Genetic diversity | Soil pH |
|---|---|---|---|---|---|---|---|---|---|---|---|---|---|---|---|
| AA016 | F | VT | 2x | 4 | 20 | 8 | 15 | SK | Košický kraj: Podlesok, rocks in the entrance to Suchá Belá gorge, limestone rocks | E. Záveská, J. Kučera, F. Kolář | 48.9603 | 20.3833 | 600 | 0.051 | 7.82 |
| AA021 | F | VT | 2x | 4 | 20 | 3 | | SK | Prešovský kraj: Tatranská kotlina, rocks next to the entrance to Belianska jaskyňa cave, slopes and limestone rocks in mixed forest | E. Záveská, F. Kolář | 49.2289 | 20.3117 | 900 | | 7.26 |
| AA023 | F | VT | 2x | 4 | 19 | 2 | | SK | Žilinský kraj: Bešeňová, travertine rock 1 km NNE from the church in the village, travertine rock | E. Záveská, F. Kolář | 49.1073 | 19.4347 | 574 | | 7.28 |
| AA042 | F | NT | 4x | 2 | 20 | 3 | | AT | Steiermark: Öblarn, slopes of valley of Walchenbach S of the village, rocky valley | F. Kolář | 47.4508 | 14.0064 | 713 | | 7.52 |
| AA065 | A | FG | 4x | 3 | 15 | 4 (BAL) | 13 | RO | Argeş: Făgăraş Mts, slopes above lake Balea, close to Transfăgăraş road, alpine scree | F. Kolář, G. Šrámková | 45.602 | 24.6226 | 2269 | | 5.58 |
| AA067 | F | FG | 4x | 3 | 20 | 4 (DRA) | 13 | RO | Argeş: Dâmbovicioara, river canyon W of the village, shady rocks in a canyon | F. Kolář, G. Šrámková | 45.4416 | 25.2239 | 858 | 0.046 | 7.65 |
| AA075 | F | RD | 4x | 1 | 15 | 4 | | RO | Suceava: Cârlibaba, rocky slope above the road to Borşa, W of the village, shady rocks, eroded slope | F. Kolář, G. Šrámková | 47.5759 | 25.0771 | 981 | 0.058 | 7.56 |
| AA084 | A | VT | 2x | 5 | 20 | 4 (VEL) | 8 | SK | Prešovský kraj: Vysoké Tatry, wet rocks and gravel in Velická Dolina ravine, between Sliezsky dom chalet and Dlhé pleso lake, alluvial gravel, wet rocks | F. Kolář, E. Záveská, S. Španiel, M. Kolník, K. Marhold | 49.162 | 20.1542 | 1823 | 0.048 | 5.23 |
| AA086 | A | VT | 2x | 4 | 12 | 2 | | SK | Žilinský kraj: Zuberec, Zverovka, limestone rocks at the top of Osobitá mountain, exposed rocks, gravel | F. Kolář, E. Záveská, S. Španiel, M. Kolník, K. Marhold | 49.2593 | 19.7218 | 1552 | | 6.11 |
| AA090 | A | VT | 2x | 5 | 17 | 4 | 11 | SK | Prešovský kraj: Vysoké Tatry, screes and rocks above the Zelené pleso lake up to Medený vodopád waterfall and Dlhý vodopád waterfall, scree, wet rocks | F. Kolář, E. Záveská, S. Španiel, M. Kolník, K. Marhold | 49.2065 | 20.2151 | 1625 | 0.048 | 5.53 |

| | | | | | | | | | | | | | | | |
|---|---|---|---|---|---|---|---|---|---|---|---|---|---|---|---|
| AA091 | A | VT | 2x | 4 | 20 | 3 | | SK | Prešovský kraj: Ždiar, Belianské Tatry, along blue tourist path in Kopské sedlo pass, exposed rocks, eroded slope | F. Kolář, E. Záveská, S. Španiel, M. Kolník, K. Marhold | 49.2299 | 20.219 | 1751 | | 7.14 |
| AA096 | F | ZT | 4x | 4 | 19 | 4 | | SK | Žilinský kraj: Nízke Tatry, Liptovský Ján, Jánska dolina valley, S of the village, alluvial gravel, eroded slope above road | F. Kolář, S. Španiel, M. Kolník, K. Marhold; J. Mořkovská | 49.0108 | 19.6733 | 744 | 0.045 | 7.15 |
| AA145 | A | NT | 4x | 2 | 19 | 3 | | AT | Steiermark: Niedere Tauern, valley and rocks along the trekking path towards Gamskögel mountain, moist valley (with Petasites sp.) and rocks | E. Záveská, S. Španiel | 47.3733 | 14.554 | 1752 | | NA |
| AA162 | A | VT | 2x | 5 | 20 | 2 | | SK | Prešovský kraj: Vysoké Tatry, Mlynická dolina valley, Skok waterfall, wet rocks, alluvial gravel | F. Kolář, M. Lučanová, K. Marhold, J. Smatanová, J. Mořkovská | 49.1534 | 20.0458 | 1750 | | 5.66 |
| AA167 | A | ZT | 4x | 4 | 16 | 3 | | SK | Žilinský kraj: Pribylina, Račkova dolina valley and Račkove plesá lakes, open places in grasslands disturbed by marmots | M. Lučanová, K. Marhold | 49.2 | 19.8047 | 1690 | | 5.16 |
| AA168 | A | ZT | 4x | 5 | 12 | 4 (TKO) | 14 | SK | Žilinský kraj: Zuberec, Zverovka, rocky slopes above Horné Roháčske pleso lake and on N slopes of Tri Kopy mountain, wet rocks, screes | F. Kolář, K. Marhold | 49.2045 | 19.7352 | 1783 | 0.052 | 4.83 |
| AA169 | A | ZT | 4x | 5 | 12 | 3 | 15 | SK | Žilinský kraj: Vysoké Tatry, northern rocky slopes under the Baníkovské sedlo saddle, wet rocks | M. Lučanová, J. Smatanová, J. Mořkovská | 49.2012 | 19.7084 | 1864 | | 5.31 |
| AA170_2 | A | VT | 2x | 4 | 28 | 4 (TRD) | | SK | Prešovský kraj: Žiar, Tristárska dolina valley, open gravely sites, screes | F. Kolář, M. Lučanová, K. Marhold, J. Smatanová, J. Mořkovská | 49.2502 | 20.2053 | 1650 | 0.051 | 6.31 |
| AA170_4 | A | ZT | 4x | 4 | 20 | 4 (TRT) | | SK | Prešovský kraj: Žiar, Tristárska dolina valley, open gravely sites, screes | F. Kolář, M. Lučanová, K. Marhold, J. Smatanová, J. Mořkovská | 49.2502 | 20.2053 | 1809 | 0.05 | 7.51 |
| AA171 | F | ZT | 4x | 4 | 19 | 4 | 15 | SK | Prešovský kraj: Hranovnica, slopes above the road to Poprad town, S of the Kvetnica village, rocky eroded slope above the road and open oak forest | F. Kolář, M. Lučanová, J. Mořkovská | 49.0072 | 20.2864 | 720 | 0.057 | 6.46 |
| AA172 | F | ZT | 4x | 4 | 20 | 3 | | SK | Prešovský kraj: Primovce, Primovské skaly rocks in the village, shady rocks | F. Kolář, M. Lučanová, J. Mořkovská | 49.0159 | 20.3825 | 605 | | 5.39 |
| AA173 | F | VT | 2x | 4 | 17 | 3 | | SK | Prešovský kraj: Gánovce, travertine outcrop in the village, exposed rocks | F. Kolář, M. Lučanová, J. Mořkovská | 49.0299 | 20.3207 | 664 | | 7.29 |

| | | | | | | | | | | | | | | | |
|---|---|---|---|---|---|---|---|---|---|---|---|---|---|---|---|
| AA176 | A | VT | 2x | 5 | 17 | 3 | | SK | Prešovský kraj: Vysoké Tatry, Mengusovská dolina valley, Veľké Hincovo pleso lake, open moist gravely patches, lake shore | F. Kolář | 49.1756 | 20.0605 | 1950 | | 6.01 |
| AA182 | F | VT | 2x | 4 | 20 | 3 | | SK | Žilinský kraj: Kráľova Lehota, rock above Čierny Váh river, exposed dry rock | K. Marhold, J. Smatanová; J. Mořkovská | 49.0153 | 19.8111 | 670 | | 6.94 |
| AA183 | F | VT | 2x | 4 | 20 | 4 | | SK | Žilinský kraj: Malužiná, slope above road to the quarry, eroded slope | K. Marhold, J. Smatanová; J. Mořkovská | 48.9845 | 19.7573 | 735 | 0.045 | 7.24 |
| AA184 | F | VT | 2x | 4 | 19 | 3 | | SK | Žilinský kraj: Liptovský Hrádok, slopes above Čierny Váh river, near Borová Sihoť spring, eroded slope | K. Marhold, J. Smatanová; J. Mořkovská | 49.0386 | 19.7005 | 624 | | 7.21 |
| AA208 | F | VT | 2x | 4 | 38 | 8 | 14 | SK | Prešovský kraj: Svit, Baba hill, SE slopes, forest clearing in pine forest | J. Mořkovská, J. Smatanová | 49.0435 | 20.1808 | 844 | 0.05 | 7.54 |
| AA220 | F | RD | 4x | NA | 12 | 0 | | RO | Bistrita-Năsăud: Parva, slopes above the road in a deep valley of Rebra brook at N margin of the village, open sites above road in deciduous forest | F. Kolář, M. Bohutínská | 47.4024 | 24.5461 | 545 | | 6.09 |
| AA222 | A | FG | 4x | 3 | 16 | 4 (LAC) | 15 | RO | Argeş: Făgărăș Mts., slopes above path from second bend of the Transfăgărășan road S of the tunnel to Lacul Capra lake, calcareous rocks and scree | F. Kolář, M. Bohutínská | 45.5954 | 24.6346 | 2092 | 0.042 | 7.5 |
| AA226 | F | ZT | 4x | 4 | 12 | 3 | | SK | Žilinský kraj: Prosiek, transect through Prosiecka dolina valley, N of the village, semi-shady rocks in a gorge | J. Mořkovská | 49.16 | 19.4962 | 656 | | 7.19 |
| AA227 | A | VT | 2x | 5 | 21 | 2 | | SK | Prešovský kraj: Vysoké Tatry, ca 300 meters E of Prielom saddle, along the blue marked tourist path, alpine scree | J. Mořkovská | 49.1762 | 20.15 | 2070 | | 6.94 |
| AA228 | A | VT | 2x | 5 | 17 | 3 | | SK | Prešovský kraj: Vysoké Tatry, north facing slope below Pod Polskym hrebeňom saddle, along tourist path, rocks | J. Mořkovská | 49.1738 | 20.1391 | 2147 | | 5.19 |
| AA229 | F | ZT | 4x | 4 | 15 | 4 | 15 | SK | Žilinský kraj: Huty, rocks in the entrance to Kvačianska dolina gorge, limestone rocks | E. Záveská, J. Kučera, F. Kolář | 49.2076 | 19.5494 | 750 | 0.038 | 7.17 |
| AA234 | F | ZT | 4x | 4 | 19 | 4 | | SK | Prešovský kraj: Lesnica, N facing slopes of the Dunajec river canyon 1.5 km NW of the village, shady rocks in beech forest | F. Kolář, G. Šrámková, J. Smatanová | 49.4117 | 20.4488 | 437 | 0.044 | 7.26 |
| AA236 | F | VT | 2x | 4 | 19 | 4 | | SK | Prešovský kraj: Lipovce, at the lowest waterfall in Kamenná Baba gorge, 1.5 km W of the village, shady limestone rocks | F. Kolář, G. Šrámková, J. Smatanová | 49.0589 | 20.9306 | 659 | 0.042 | 6.82 |
| AA242 | F | ZT | 4x | 4 | 20 | 3 | | SK | Košiský kraj: Spišká Nová Ves - Čingov, N facing slopes above Hornád river, 500 m WSW of the settlement, shady north-facing rock | J. Mořkovská, K. Marhold, S. Španiel, J. Smatanová | 48.9402 | 20.4774 | 530 | | 7.29 |

| | | | | | | | | | | | | | | |
|---|---|---|---|---|---|---|---|---|---|---|---|---|---|---|
| AA244 | F | VT | 2x | 4 | 12 | 3 | | SK | Žilinský kraj: Šútovo, small limestone hill at E margin of the village, rocky and gravely sunny slope | F. Kolář, A. Knotek, G. Šrámková | 49.15197 | 19.085165 | 480 | | 7.41 |
| AA248 | A | VT | 2x | 5 | 4 | 4 | | SK | Prešovský kraj: Rysy mountain, just below the summit, rocky outcrop near the summit | M. Bohutínská | 49.1795 | 20.0881 | 2488 | 0.041 | 5.43 |
| AA251 | F | FG | 4x | 3 | 14 | 3 | 15 | RO | Argeş: Dâmbovicioara, river canyon S of the village, gravel along the road, most likely secondary habitat, not on the rocks only below them in the gravel | E. Záveská | 45.4267 | 25.2133 | 915 | | 8.3 |
| AA252 | F | NT | 4x | 2 | 19 | 3 | 15 | AT | Steiermark: Schönberg-Lachtal, along a road L 514, ca 600 m N of of Hocheggerstraße and Glattjoch Straße junction, gravel and stones along the road | F. Kolář, M. Hanzl, E. Záveská, S. Španiel | 47.1826 | 14.3379 | 856 | | 8.02 |
| AA253 | A | NT | 4x | 2 | 21 | 4 (SCH) | 15 | AT | Steiermark: Niedere Tauern, Schießeck mountain, N exposed rocky slopes, screes and rocks, quartzite | F. Kolář, M. Hanzl, S. Španiel | 47.2777 | 14.3219 | 2240 | 0.033 | 6.7 |
| AA254 | A | NT | 4X | NA | 0 | 0 | 15 | AT | Steiermark: Seckauer Alpen: Hochreichart, northwestern crest, stabilized amphibolite screes | P. Schönswetter | 47.36444 | 14.68083 | 2360 | | NA |
| AA255 | F | NT | 4x | 2 | 20 | 4 | 15 | AT | Steiermark: Seckauer Alpen, lower-most Ingeringgraben, 1.6 km NNW of Wasserberg castle, Ingeringbach valley, siliceous rocks | P. Schönswetter | 47.2842 | 14.6819 | 970 | 0.052 | 4.72 |
| AA293 | A | RD | 4x | 1 | 4 | 3 | | UA | Zakarpatská oblast: Dragobrat, bellow the top of Blyznytsya mountain, S slope, in glacial cirque, 70 - 100 vertical meters above Ivor lakes, limestone in glacial cirque | M. Bohutínská, J. Hojka, D. Bohutínský, F. Rooks | 48.2286 | 24.2323 | 1614 | | 5.44 |
| AA294 | A | RD | 4x | 1 | 4 | 3 | | RO | Maramureş: Borșa, around the entrance to glacial cirque at E slopes of Pietrosul Rodnei mountain, along the dirt road, 5.7 km SSW of the town, rocky slope with sparse vegetation | F. Kolář, M. Bohutínská | 47.6031 | 24.6488 | 1780 | | 6.98 |
| AA295 | A | RD | 4x | 1 | 12 | 3 | | UA | Zakarpatská oblast: Svidovec, Gerisaska mountain, upper part of the glacial cirque (below the rocks) on the eastern slope, rocks and scree in glacial cirque | J. Chrtek, K. Kabátová | 48.2722 | 24.1623 | 1702 | | 6.38 |
| AA296 | A | RD | 4x | 1 | 10 | 3 | | RO | Maramureş: Borșa, rocks at the NW facing slopes of glacial cirque at E slopes of Pietrosul Rodnei mountain, 6 km SSW of the town, rocky crevices | F. Kolář, M. Bohutínská | 47.5977 | 24.6469 | 1820 | | 5.15 |
| AA298 | A | RD | 4x | 1 | 18 | 4 | | RO | Maramureş: Borșa, NW slopes of Vârful Ciungilor mountain, in Bila stream and on the rock above spring of the stream, limestone | M. Bohutínská, J. Hojka, D. Bohutínský, F. Rooks | 47.5393 | 24.8868 | 1586 | 0.06 | 6.63 |
| AA299 | A | RD | 4x | 1 | 10 | 2 | | RO | Maramureş: Borșa, NW slope of the NW glacial cirque bellow Ineu peak, glacial cirque | M. Bohutínská, J. Hojka, D. Bohutínský, F. Rooks | 47.5273 | 24.8806 | 2017 | | 7.12 |

| | | | | | | | | | | | | | | |
|---|---|---|---|---|---|---|---|---|---|---|---|---|---|---|
| AA300 | A | NT | 4x | 2 | 18 | 4 | | AT | Steiermark: Pusterwald, shores of Wildsee lake below Eiskarspitz mountain, snowbed in a glacial cirque | F. Kolář, A. Knotek, S. Španiel, P. Schönswetter, K. Hülber | 47.3256 | 14.2304 | 2117 | 0.056 | 6.02 |
| AA302 | A | NT | 4x | 2 | 17 | 3 | | AT | Steiermark: Krakaudorf, screes in Sauoffensee glacial cirque, W of the lake, wet rocks and scree slope below | F. Kolář, P. Schönswetter, K. Hülber | 47.2586 | 14.0103 | 2184 | | 6.16 |
| AA303 | A | NT | 4x | 2 | 20 | 2 | | AT | Steiermark: Krakaudorf, slope above Sauoffensee lake, stony snowbed slope | A. Knotek, S. Španiel | 47.2588 | 14.0049 | 2030 | | 5.54 |
| AA304 | F | NT | 4x | 2 | 20 | 3 | | AT | Steiermark: Aigen, rocks in Gullingtal valley, next to the bridge W of a quarry, rocks | F. Kolář, A. Knotek, S. Španiel | 47.4936 | 14.1722 | 800 | | 7.74 |
| AA322 | A | ZT | 4x | 4 | 20 | 3 | | SK | Nizke Tatry: NE slope between Krakova hola mountain and Pusté mountain, rocky sites in forest clearing | J. Mořkovská | 48.9919 | 19.623 | 1472 | | 7.07 |
| AA330 | F | FG | 4x | 3 | 13 | 4 | | RO | Brasov: Timisu de Sus, rocks above railway, ca 5 km NNE of the village, shady rocks, travertine spring | F. Kolář, M. Bohutínská, D. Požárová, F. Rooks | 45.57 | 25.6083 | 797 | 0.063 | 8 |
| AA335 | F | FG | 4x | 3 | 20 | 3 | | RO | Brasov: Zărneşti, narrowest part of Prăpăstiile Zărneştilor gorge, 5.5 km SW of the railway station in the town, rocks in the gorge | F. Kolář, M. Bohutínská, D. Požárová, F. Rooks | 45.5253 | 25.2812 | 978 | | 6.76 |
| AA338 | F | NT | 4x | 2 | 20 | 4 (HOC) | | AT | Steiermark: Mautstatt, Calcareous rocks and screes near road and railroad, ca 4 km NE from the village, calcareous rocks and bank between railroad and small river | A. Knotek; D. Požárová | 47.37 | 15.3867 | 560 | 0.044 | 7.61 |
| AA339 | F | NT | 4x | 2 | 18 | 4 (KAS) | | AT | Kärnten: Sankt Paul im Lavanttal , castle ruin Rabenstein, circa 2 km S from the town, rocks and walls of castle ruins | A. Knotek; D. Požárová | 46.6883 | 14.8717 | 620 | 0.045 | 7.98 |
| AA340 | F | NT | 4x | 2 | 20 | 4 (KOS) | | AT | Oberösterreich: Mitterweißenbach, Kesselbach stream riverbank near the road from Bad Ischl to Ebensee, calcareous river bank | A. Knotek; D. Požárová | 47.7469 | 13.6897 | 460 | 0.048 | 8.42 |

**Table S2** Details on diversity and accession codes of genome resequenced populations used for coalescent simulations

| pop code | Region | Ecotype | N indivs | N all sites called | N SNPs | Genetic diversity (pi) | original code | BioSample codes (SRA) |
|---|---|---|---|---|---|---|---|---|
| AA300 | NT | Alpine | 8 | 3939154 | 361293 | 0.0267 | WIL | SAMN13420543-50 |
| AA339 | NT | Foothill | 8 | 2920550 | 262141 | 0.0255 | KOS | SAMN09759803-09 |
| AA065 | FG | Alpine | 8 | 3950174 | 349273 | 0.0260 | BAL | SAMN13420559-66 |
| AA067 | FG | Foothill | 8 | 2218553 | 176496 | 0.0213 | DRA | SAMN09759740-47 |
| AA299 | RD | Alpine | 8 | 3925916 | 417156 | 0.0286 | INE | SAMN13420575-81 |
| AA075 | RD | Foothill | 8 | 3917955 | 409493 | 0.0213 | CAR | SAMN13420582-89 |
| AA168 | ZT | Alpine | 8 | 3723199 | 354763 | 0.0285 | TKO | SAMN09759921-28 |
| AA171 | ZT | Foothill | 8 | 3918890 | 400976 | 0.0275 | HRA | SAMN13420590-97 |
| AA084 | VT | Alpine | 8 | 3447202 | 239213 | 0.0234 | VEL | SAMN09759960-67 |
| AA016 | VT | Foothill | 8 | 3915391 | 317662 | 0.0267 | SUB | SAMN13420527-34 |

**Table S3** Pearson pairwise correlation coefficients between the environmental parameters (upper) and morphological characters (lower) used in the analyses.

| Ecology | PAR | Precipitation | Temperature | EIV_Moisture | EIV_Nutrients | soil_ph | EIV_Light | Veget_ cover |
|---|---|---|---|---|---|---|---|---|
| **PAR** | | -0.69 | 0.54 | -0.39 | 0.15 | 0.29 | -0.15 | 0.14 |
| **Precipitation** | -0.69 | | -0.65 | 0.20 | -0.45 | -0.40 | 0.47 | -0.26 |
| **Temperature** | 0.54 | -0.65 | | -0.33 | 0.54 | 0.61 | -0.56 | 0.26 |
| **EIV_Moisture** | -0.39 | 0.20 | -0.33 | | 0.44 | -0.10 | -0.28 | -0.13 |
| **EIV_Nutrients** | 0.15 | -0.45 | 0.54 | 0.44 | | 0.44 | -0.78 | 0.18 |
| **soil_ph** | 0.29 | -0.40 | 0.61 | -0.10 | 0.44 | | -0.34 | 0.17 |
| **EIV_Light** | -0.15 | 0.47 | -0.56 | -0.28 | -0.78 | -0.34 | | -0.12 |
| **Vegetation cover** | 0.14 | -0.26 | 0.26 | -0.13 | 0.18 | 0.17 | -0.12 | |

| Morphology | HP | HF | PL | PW | PL.PW | SL | SW | SL.SW | SLL | SLW | SLL.SLW | RLL | RLW | RLL.RLW | SNLP | RNLP |
|---|---|---|---|---|---|---|---|---|---|---|---|---|---|---|---|---|
| **HP** | | 0.69 | -0.21 | -0.28 | 0.17 | -0.25 | -0.36 | 0.24 | 0.50 | 0.43 | 0.05 | 0.41 | 0.37 | 0.12 | 0.41 | 0.30 |
| **HF** | 0.69 | | -0.24 | -0.26 | 0.11 | -0.18 | -0.21 | 0.10 | 0.37 | 0.26 | 0.06 | 0.33 | 0.23 | 0.18 | 0.28 | 0.17 |
| **PL** | -0.21 | -0.24 | | 0.80 | -0.03 | 0.72 | 0.55 | -0.02 | 0.09 | 0.10 | -0.04 | 0.13 | 0.15 | -0.03 | -0.03 | 0.01 |
| **PW** | -0.28 | -0.26 | 0.80 | | -0.55 | 0.64 | 0.62 | -0.19 | 0.01 | 0.03 | -0.07 | 0.06 | 0.09 | -0.04 | -0.09 | -0.03 |
| **PL.PW** | 0.17 | 0.11 | -0.03 | -0.55 | | -0.07 | -0.28 | 0.30 | 0.08 | 0.05 | 0.09 | 0.07 | 0.05 | 0.01 | 0.08 | 0.06 |
| **SL** | -0.25 | -0.18 | 0.72 | 0.64 | -0.07 | | 0.71 | 0.03 | 0.05 | 0.07 | -0.04 | 0.13 | 0.15 | -0.02 | -0.02 | 0.00 |
| **SW** | -0.36 | -0.21 | 0.55 | 0.62 | -0.28 | 0.71 | | -0.64 | -0.05 | -0.06 | -0.06 | 0.03 | 0.03 | 0.00 | -0.11 | -0.10 |
| **SL.SW** | 0.24 | 0.10 | -0.02 | -0.19 | 0.30 | 0.03 | -0.64 | | 0.13 | 0.16 | 0.03 | 0.09 | 0.11 | -0.02 | 0.14 | 0.16 |
| **SLL** | 0.50 | 0.37 | 0.09 | 0.01 | 0.08 | 0.05 | -0.05 | 0.13 | | 0.80 | 0.08 | 0.64 | 0.67 | 0.03 | 0.43 | 0.27 |
| **SLW** | 0.43 | 0.26 | 0.10 | 0.03 | 0.05 | 0.07 | -0.06 | 0.16 | 0.80 | | -0.40 | 0.53 | 0.58 | -0.02 | 0.56 | 0.29 |
| **SLL.SLW** | 0.05 | 0.06 | -0.04 | -0.07 | 0.09 | -0.04 | -0.06 | 0.03 | 0.08 | -0.40 | | 0.03 | -0.03 | 0.10 | -0.31 | -0.05 |
| **RLL** | 0.41 | 0.33 | 0.13 | 0.06 | 0.07 | 0.13 | 0.03 | 0.09 | 0.64 | 0.53 | 0.03 | | 0.77 | 0.39 | 0.25 | 0.26 |
| **RLW** | 0.37 | 0.23 | 0.15 | 0.09 | 0.05 | 0.15 | 0.03 | 0.11 | 0.67 | 0.58 | -0.03 | 0.77 | | -0.20 | 0.28 | 0.25 |
| **RLL.RLW** | 0.12 | 0.18 | -0.03 | -0.04 | 0.01 | -0.02 | 0.00 | -0.02 | 0.03 | -0.02 | 0.10 | 0.39 | -0.20 | | -0.05 | -0.01 |
| **SNLP** | 0.41 | 0.28 | -0.03 | -0.09 | 0.08 | -0.02 | -0.11 | 0.14 | 0.43 | 0.56 | -0.31 | 0.25 | 0.28 | -0.05 | | 0.46 |
| **RNLP** | 0.30 | 0.17 | 0.01 | -0.03 | 0.06 | 0.00 | -0.10 | 0.16 | 0.27 | 0.29 | -0.05 | 0.26 | 0.25 | -0.01 | 0.46 | |

**Table S4** Average values of traits in foothill and alpine groups of *A. arenosa* plants sampled in field and in common garden experiment. The measurements are provided in a Supplementary data file S3.

| | | | | Field data | | Common garden data | |
|---|---|---|---|---|---|---|---|
| Traits | Description | Category | Unit | Alpine | Foothill | Alpine | Foothill |
| HP | Height of the main stem | stem | mm | 125.05 | 269 | 114 | 185 |
| HF | Distance from the rosette to the lowest flower | stem | mm | 79.73 | 130.57 | 64.3 | 117.1 |
| PL | Petal length | flower | mm | 6.465 | 5.653 | 6.52 | 5.434 |
| PW | Petal width | flower | mm | 3.475 | 2.815 | 3.55 | 2.668 |
| PL.PW | Ratio Petal length / width | flower | mm | 1.887 | 2.047 | 1.85 | 2.09 |
| SL | Sepal length | flower | mm | 2.796 | 2.404 | 2.68 | 2.441 |
| SW | Sepal width | flower | mm | 1.428 | 1.093 | 1.14 | 1.067 |
| SL.SW | Ratio Sepal length / width | flower | mm | 2.014 | 2.264 | 2.41 | 2.332 |
| SLL | Stem leaf length | leaf | mm | 19.96 | 26.49 | 14.6 | 14.606 |
| SLW | Stem leaf width | leaf | mm | 5.897 | 7.649 | 4.57 | 5.261 |
| SLL.SLW | Ratio Stem length / width | leaf | mm | 3.721 | 3.943 | 3.35 | 2.944 |
| RLL | Rosette leaf length | leaf | mm | 31.65 | 34.36 | 26.6 | 23.401 |
| RLW | Rosette leaf width | leaf | mm | 7.385 | 8.647 | 5.58 | 5.405 |
| RLL.RLW | Ratio Rosette length / width | leaf | mm | 4.43 | 4.082 | 4.86 | 4.511 |
| SNLP | Stem leaf number of lobe pairs | leaf lobes | count | 1.865 | 2.585 | 1.6 | 2.914 |
| RNLP | Rosette leaf number of lobe pairs | leaf lobes | count | 3.488 | 4.484 | 4.81 | 6.862 |

**Table S5** Genetic and morphological diversity and differentiation of foothill and alpine populations of *A. arenosa* categorized according to four regions (treating VT and ZT regions together).

| | Genetic data | | | Morphological data from field | | | Morphological data from common garden | | |
|---|---|---|---|---|---|---|---|---|---|
| **Grouping** | **N[1]** | **AMOVA (%)[2]** | **Gen. diversity (π)[3]** | **N[1]** | **Classif.[4]** | **Differentiation[5]** | **N[1]** | **Classif.[4]** | **Differentiation[5]** |
| **Grouping by region (4 regions)** | | | | | | | | | |
| all pops. | 200 | 19 | 0.048 | 999 | 48 | 6.81*** (2.01%) | 223 | 58 | 5.04** (4.38%) |
| Foothill | 109 | 21 | 0.049 | 559 | 70 | 16.58*** (8.23%) | 117 | 68 | 9.00*** (13.6%) |
| Alpine | 91 | 28 | 0.048 | 440 | 53 | 13.97*** (8.77%) | 106 | 73 | 17.2*** (25%) |
| **Grouping by ecotype** | | | | | | | | | |
| VT+ZT | 115 | 7 | 0.047 / 0.049 | 584 | 88 | 30.41*** (4.97%) | 107 | 94 | 53.7*** (33.8%) |

[1] N individuals
[2] Among-group genetic variation component (in %) as explained by hierarchical AMOVA
[3] Pairwise nucleotide diversity (π) averaged over populations with ≥ 4 individuals (foothill / alpine ecotypes, respectively)
[4] % of correct classification into ecotype / regional group as inferred by classificatory discriminant analysis of the 16 morphological characters
[5] F-values and significance (*P < 0.05, **P < 0.01, ***P < 0.001) of permanova analysis of the 16 morphological characters

**Table S6** Analysis of molecular variance (AMOVA) reflecting regional and ecotypic clustering of *A. arenosa* in total and within each region separately.

| Grouping | dataset | SSD | MSD | % explained variance | d.f. | p |
|---|---|---|---|---|---|---|
| **Grouping by region (5 regions: NT, VT, ZT, RD, FG)** | | | | | | |
| Regional group | all pops | 0.0177 | 0.0044 | 19.60 | 4 | **< 0.001** |
| Populations within regional group | all pops | 0.0359 | 0.0007 | 39.74 | 52 | **< 0.001** |
| Individuals within population | all pops | 0.0367 | 0.0003 | 40.66 | 143 | |
| Total | all pops | 0.0904 | 0.0005 | 100.00 | 199 | |
| Regional group | foothill pops | 0.0111 | 0.0028 | 22.89 | 4 | **< 0.001** |
| Populations within regional group | foothill pops | 0.0156 | 0.0007 | 32.10 | 24 | **< 0.001** |
| Individuals within population | foothill pops | 0.0219 | 0.0003 | 45.01 | 80 | |
| Total | foothill pops | 0.0486 | 0.0005 | 100.00 | 108 | |
| Regional group | alpine pops | 0.0116 | 0.0029 | 29.80 | 4 | **< 0.001** |
| Populations within regional group | alpine pops | 0.0125 | 0.0005 | 32.04 | 23 | **< 0.001** |
| Individuals within population | alpine pops | 0.0148 | 0.0002 | 38.16 | 63 | |
| Total | alpine pops | 0.0389 | 0.0004 | 100.00 | 90 | |
| **Grouping by region (4 regions: NT, VT+ZT, RD, FG)** | | | | | | |
| Regional group | all pops | 0.0174 | 0.0058 | 19.24 | 3 | **< 0.001** |
| Populations within regional group | all pops | 0.0362 | 0.0007 | 40.1 | 53 | **< 0.001** |
| Individuals within population | all pops | 0.0367 | 0.0003 | 40.66 | 143 | |
| Total | all pops | 0.0904 | 0.0005 | 100 | 199 | |
| Regional group | foothill pops | 0.0102 | 0.0034 | **20.87** | 3 | **< 0.001** |
| Populations within regional group | foothill pops | 0.0166 | 0.0007 | 34.12 | 25 | **< 0.001** |
| Individuals within population | foothill pops | 0.0219 | 0.0003 | 45.01 | 80 | |
| Total | foothill pops | 0.0486 | 0.0005 | 100 | 108 | |
| Regional group | alpine pops | 0.0108 | 0.0036 | **27.9** | 3 | **< 0.001** |
| Populations within regional group | alpine pops | 0.0132 | 0.0005 | 33.94 | 24 | **< 0.001** |
| Individuals within population | alpine pops | 0.0148 | 0.0002 | 38.16 | 63 | |
| Total | alpine pops | 0.0389 | 0.0004 | 100 | 90 | |
| **Grouping by ecotype** | | | | | | |
| Elevational group | all pops | 0.0028 | 0.0028 | 3.12 | 1 | **< 0.001** |
| Populations within elevational group | all pops | 0.0508 | 0.0009 | 56.22 | 55 | **< 0.001** |
| Individuals within population | all pops | 0.0367 | 0.0003 | 40.66 | 143 | |
| Total | all pops | 0.0904 | 0.0005 | 100 | 199 | |
| Elevational group | NT region | 0.0006 | 0.0006 | 4.87 | 1 | 0.085 |
| Populations within elevational group | NT region | 0.0049 | 0.0005 | 42 | 10 | **< 0.001** |
| Individuals within population | NT region | 0.0062 | 0.0002 | 53.13 | 29 | |
| Total | NT region | 0.0117 | 0.0003 | 100 | 40 | |
| Elevational group | VT + ZT region | 0.0036 | 0.0036 | 7.39 | 1 | **< 0.001** |
| Populations within elevational group | VT + ZT region | 0.0156 | 0.0005 | 31.62 | 30 | **< 0.001** |
| Individuals within population | VT + ZT region | 0.03 | 0.0004 | 60.99 | 83 | |
| Total | VT + ZT region | 0.0493 | 0.0004 | 100 | 114 | |
| Elevational group | VT region | 0.0011 | 0.0011 | 2.77 | 1 | 0.7393 |
| Populations within elevational group | VT region | 0.0178 | 0.001 | 45.86 | 17 | **< 0.001** |
| Individuals within population | VT region | 0.0199 | 0.0004 | 51.37 | 52 | |

| | | | | | | |
|---|---|---|---|---|---|---|
| Total | VT region | 0.0387 | 0.0006 | 100 | 70 | |
| Elevational group | ZT region | 0.0003 | 0.0003 | 3.39 | 1 | 0.7972 |
| Populations within elevational group | ZT region | 0.0041 | 0.0004 | 40.54 | 11 | < **0.001** |
| Individuals within population | ZT region | 0.0057 | 0.0002 | 56.07 | 31 | |
| Total | ZT region | 0.0102 | 0.0002 | 100 | 43 | |
| Elevational group | RD region | 0.0004 | 0.0004 | 6.35 | 1 | 0.571 |
| Populations within elevational group | RD region | 0.002 | 0.0004 | 34.94 | 5 | < **0.001** |
| Individuals within population | RD region | 0.0034 | 0.0002 | 58.71 | 15 | |
| Total | RD region | 0.0058 | 0.0003 | 100 | 21 | |
| Elevational group | FG region | 0.0006 | 0.0006 | 9.19 | 1 | 0.121 |
| Populations within elevational group | FG region | 0.0016 | 0.0004 | 24.97 | 4 | **0.043** |
| Individuals within population | FG region | 0.0041 | 0.0003 | 65.84 | 16 | |
| Total | FG region | 0.0062 | 0.0003 | 100 | 21 | |

**Table S7** Genetic and morphological diversity and differentiation of foothill and alpine populations of *A. arenosa* from the five sampled mountain regions.

| | Genetic diversity and differentiation | | | | Field morphology | | | Common garden morphology | | |
|---|---|---|---|---|---|---|---|---|---|---|
| | N[1] | AMOVA (%)[2] | Genetic diversity [3] | Pairwise Fst[4] | N[1] | CDA[5] (%) | Differentiation[6] | N[1] | CDA[5] (%) | Differentiation[6] |
| **Grouping by region** | | | | | | | | | | |
| All populations | 200 | 20 | 0.048 | 0.132 | 999 | 43 | $6.43^{***}$ | 223 | 47 | $4.00^{*}$ |
| Foothill | 109 | 23 | 0.049 | 0.120 | 559 | 65 | $14.1^{***}$ | 117 | 62 | $6.58^{***}$ |
| Alpine | 91 | 30 | 0.048 | 0.139 | 440 | 48 | $10.75^{***}$ | 106 | 75 | $19.94^{***}$ |
| **Grouping by ecotype** | | | | | | | | | | |
| All regions | 200 | 3 | 0.049/0.048 | | 999 | 89 | $49.25^{***}$ | 223 | 83 | $108.5^{***}$ |
| NT (Niedere Tauern) | 41 | n.s. | 0.047/0.045 | 0.105/0.089 | 232 | 100 | $8.83^{**}$ | 60 | 63 | $6.52^{**}$ |
| VT (Vysoké Tatry) | 73 | n.s. | 0.047/0.047 | 0.066/0.068 | 380 | 89 | $17.15^{***}$ | 48 | 83 | $60.91^{***}$ |
| ZT (Západné Tatry) | 42 | n.s. | 0.046/0.051 | 0.049/0.045 | 204 | 90 | $16.36^{***}$ | 59 | 93 | $17.00^{***}$ |
| RD (Rodna) | 22 | n.s. | 0.058/0.060 | -/- | 85 | 93 | $3.42^{*}$ | - | - | - |
| FG (Făgăraș) | 22 | n.s. | 0.055/0.042 | 0.062/0.119 | 98 | 100 | $50.96^{***}$ | 56 | 95 | $113.61^{***}$ |

[1] N individuals
[2] Among-group genetic variation component (in %) as explained by hierarchical AMOVA
[3] Pairwise nucleotide diversity ($\pi$) averaged over populations with ≥ 4 individuals (foothill / alpine ecotypes, respectively)
[4] Pairwise Fst averaged over populations with ≥ 4 individuals (foothill / alpine ecotypes, respectively)
[5] % of correct classification into ecotype / regional group as inferred by classificatory discriminant analysis of the 16 morphological characters
[6] F-values and significance (*P < 0.05, **P < 0.01, ***P < 0.001) of permanova analysis of the 16 morphological characters

**Table S8** Pairwise differentiation (Fst) calculated based on all RAD-sequenced populations with > 4 individuals.

| | ecotype | region | ploidy | AA016 | AA065 | AA067 | AA075 | AA084 | AA090 | AA096 | AA168 | AA170_2 | AA170_4 | AA171 | AA183 | AA208 | AA222 | AA229 | AA234 | AA236 | AA248 | AA253 | AA255 | AA298 | AA300 | AA330 | AA338 | AA339 | AA340 |
|---|---|---|---|---|---|---|---|---|---|---|---|---|---|---|---|---|---|---|---|---|---|---|---|---|---|---|---|---|---|
| AA016 | F | VT | 2x | NA | | | | | | | | | | | | | | | | | | | | | | | | | |
| AA065 | A | FG | 4x | 0.16 | NA | | | | | | | | | | | | | | | | | | | | | | | | |
| AA067 | F | FG | 4x | 0.17 | 0.12 | NA | | | | | | | | | | | | | | | | | | | | | | | |
| AA075 | F | RD | 4x | 0.15 | 0.14 | 0.13 | NA | | | | | | | | | | | | | | | | | | | | | | |
| AA084 | A | VT | 2x | 0.13 | 0.16 | 0.18 | 0.17 | NA | | | | | | | | | | | | | | | | | | | | | |
| AA090 | A | VT | 2x | 0.14 | 0.17 | 0.19 | 0.17 | 0.03 | NA | | | | | | | | | | | | | | | | | | | | |
| AA096 | F | ZT | 4x | 0.07 | 0.14 | 0.15 | 0.14 | 0.1 | 0.1 | NA | | | | | | | | | | | | | | | | | | | |
| AA168 | A | ZT | 4x | 0.08 | 0.13 | 0.15 | 0.14 | 0.05 | 0.06 | 0.06 | NA | | | | | | | | | | | | | | | | | | |
| AA170_2 | A | VT | 2x | 0.08 | 0.15 | 0.17 | 0.15 | 0.1 | 0.11 | 0.08 | 0.07 | NA | | | | | | | | | | | | | | | | | |
| AA170_4 | A | ZT | 4x | 0.08 | 0.13 | 0.15 | 0.13 | 0.07 | 0.08 | 0.06 | 0.04 | 0.05 | NA | | | | | | | | | | | | | | | | |
| AA171 | F | ZT | 4x | 0.06 | 0.12 | 0.13 | 0.12 | 0.08 | 0.08 | 0.05 | 0.04 | 0.07 | 0.04 | NA | | | | | | | | | | | | | | | |
| AA183 | F | VT | 2x | 0.06 | 0.17 | 0.18 | 0.16 | 0.14 | 0.15 | 0.06 | 0.09 | 0.09 | 0.09 | 0.08 | NA | | | | | | | | | | | | | | |
| AA208 | F | VT | 2x | 0.04 | 0.16 | 0.16 | 0.14 | 0.12 | 0.12 | 0.06 | 0.07 | 0.07 | 0.06 | 0.05 | 0.05 | NA | | | | | | | | | | | | | |
| AA222 | A | FG | 4x | 0.21 | 0.12 | 0.17 | 0.19 | 0.22 | 0.23 | 0.19 | 0.18 | 0.21 | 0.18 | 0.18 | 0.22 | 0.21 | NA | | | | | | | | | | | | |
| AA229 | F | ZT | 4x | 0.08 | 0.15 | 0.16 | 0.15 | 0.11 | 0.11 | 0.05 | 0.07 | 0.09 | 0.07 | 0.06 | 0.08 | 0.07 | 0.2 | NA | | | | | | | | | | | |
| AA234 | F | ZT | 4x | 0.08 | 0.15 | 0.16 | 0.14 | 0.09 | 0.1 | 0.04 | 0.06 | 0.08 | 0.06 | 0.05 | 0.08 | 0.07 | 0.2 | 0.05 | NA | | | | | | | | | | |
| AA236 | F | VT | 2x | 0.08 | 0.2 | 0.21 | 0.18 | 0.16 | 0.16 | 0.09 | 0.12 | 0.11 | 0.11 | 0.1 | 0.09 | 0.07 | 0.25 | 0.11 | 0.1 | NA | | | | | | | | | |
| AA248 | A | VT | 2x | 0.13 | 0.18 | 0.2 | 0.18 | 0.04 | 0.05 | 0.09 | 0.07 | 0.11 | 0.09 | 0.09 | 0.14 | 0.12 | 0.24 | 0.09 | 0.08 | 0.16 | NA | | | | | | | | |
| AA253 | A | NT | 4x | 0.18 | 0.2 | 0.2 | 0.19 | 0.18 | 0.2 | 0.15 | 0.15 | 0.17 | 0.15 | 0.14 | 0.19 | 0.17 | 0.25 | 0.16 | 0.16 | 0.22 | 0.2 | NA | | | | | | | |
| AA255 | F | NT | 4x | 0.15 | 0.17 | 0.18 | 0.16 | 0.15 | 0.16 | 0.13 | 0.12 | 0.14 | 0.12 | 0.11 | 0.16 | 0.14 | 0.22 | 0.13 | 0.13 | 0.18 | 0.17 | 0.12 | NA | | | | | | |
| AA298 | A | RD | 4x | 0.17 | 0.14 | 0.14 | 0.07 | 0.18 | 0.19 | 0.15 | 0.15 | 0.17 | 0.15 | 0.13 | 0.18 | 0.16 | 0.2 | 0.16 | 0.16 | 0.2 | 0.2 | 0.2 | 0.17 | NA | | | | | |
| AA300 | A | NT | 4x | 0.12 | 0.14 | 0.15 | 0.14 | 0.13 | 0.14 | 0.1 | 0.09 | 0.12 | 0.09 | 0.09 | 0.14 | 0.12 | 0.2 | 0.11 | 0.11 | 0.16 | 0.14 | 0.09 | 0.08 | 0.15 | NA | | | | |
| AA330 | F | FG | 4x | 0.14 | 0.1 | 0.06 | 0.11 | 0.15 | 0.16 | 0.13 | 0.12 | 0.14 | 0.12 | 0.11 | 0.16 | 0.14 | 0.15 | 0.14 | 0.13 | 0.18 | 0.17 | 0.18 | 0.15 | 0.12 | 0.13 | NA | | | |
| AA338 | F | NT | 4x | 0.13 | 0.15 | 0.15 | 0.13 | 0.15 | 0.15 | 0.11 | 0.11 | 0.13 | 0.11 | 0.1 | 0.15 | 0.13 | 0.2 | 0.12 | 0.12 | 0.17 | 0.16 | 0.14 | 0.11 | 0.15 | 0.09 | 0.13 | NA | | |
| AA339 | F | NT | 4x | 0.16 | 0.17 | 0.18 | 0.16 | 0.16 | 0.17 | 0.14 | 0.13 | 0.15 | 0.13 | 0.12 | 0.17 | 0.15 | 0.23 | 0.15 | 0.14 | 0.19 | 0.18 | 0.16 | 0.13 | 0.17 | 0.1 | 0.15 | 0.12 | NA | |
| AA340 | F | NT | 4x | 0.12 | 0.15 | 0.15 | 0.14 | 0.13 | 0.14 | 0.1 | 0.1 | 0.12 | 0.1 | 0.09 | 0.13 | 0.12 | 0.2 | 0.11 | 0.1 | 0.16 | 0.14 | 0.11 | 0.08 | 0.15 | 0.06 | 0.13 | 0.09 | 0.1 | NA |

**Table S9** Parallel and non-parallel morphological differentiation in 16 traits in foothill vs. alpine populations of *A. arenosa* based on field-collected samples and plants cultivated in common garden quantified using generalized linear model, calculated separately for each trait sampled in field and in common garden experiment.

| | | Field data | | | | | | Common garden data | | | | | |
|---|---|---|---|---|---|---|---|---|---|---|---|---|---|
| | | Test statistics[1] | | | Effect sizes[2] | | | Test statistics[1] | | | Effect sizes [2] | | |
| Trait | Category | Ecotype[1] | Region[1] | E:R[1] | Ecotype | Region | E × R | Ecotype[1] | Region | E:R[1] | Ecotype | Region | E × R |
| HP | stem | 151.74$^{***}$ | 3.25$^{*}$ | 2.02 | 0.48 | 0.06 | 0.05 | 47.06$^{***}$ | 2.62 | 3.36 | 0.43 | 0.11 | 0.15 |
| HF | stem | 25.98$^{***}$ | 11.90$^{***}$ | 3.18$^{*}$ | 0.16 | 0.23 | 0.08 | 54.14$^{***}$ | 1.61 | 7.05$^{*}$ | 0.52 | 0.08 | 0.30 |
| PL | flower | 10.60$^{**}$ | 1.54 | 0.59 | 0.11 | 0.07 | 0.02 | 16.33$^{**}$ | 0.77 | 2.25 | 0.27 | 0.06 | 0.13 |
| PW | flower | 23.62$^{***}$ | 2.75$^{*}$ | 0.33 | 0.20 | 0.09 | 0.01 | 19.43$^{**}$ | 1.08 | 0.75 | 0.37 | 0.10 | 0.06 |
| PL.PW | flower | 21.79$^{***}$ | 1.8 | 1.11 | 0.09 | 0.02 | 0.01 | 6.63$^{*}$ | 0.91 | 0.22 | 0.16 | 0.07 | 0.02 |
| SL | flower | 17.38$^{***}$ | 1.04 | 0.33 | 0.15 | 0.03 | 0.01 | 13.97$^{**}$ | 3.75 | 5.26$^{*}$ | 0.09 | 0.08 | 0.10 |
| SW | flower | 55.49$^{***}$ | 3.40$^{*}$ | 0.63 | 0.27 | 0.06 | 0.01 | 6.10$^{*}$ | 1.28 | 0.22 | 0.03 | 0.02 | 0.11 |
| SL.SW | flower | 36.75$^{***}$ | 4.52$^{*}$ | 1.69 | 0.10 | 0.05 | 0.02 | 1.96 | 1.25 | 1.78 | 0.01 | 0.02 | 0.02 |
| SLL | leaf | 11.11$^{**}$ | 0.55 | 4.92$^{**}$ | 0.05 | 0.01 | 0.10 | 0.01 | 1.83 | 3.66 | 0.00 | 0.14 | 0.21 |
| SLW | leaf | 5.90$^{*}$ | 1.01 | 10.95$^{***}$ | 0.02 | 0.02 | 0.19 | 3.7 | 3.69 | 6.91$^{*}$ | 0.07 | 0.18 | 0.25 |
| SLL.SLW | leaf | 1.49 | 0.57 | 6.83$^{***}$ | 0.00 | 0.01 | 0.09 | 4.69 | 0.24 | 0.36 | 0.06 | 0.01 | 0.02 |
| RLL | leaf | 0.86 | 0.87 | 2.23 | 0.00 | 0.02 | 0.05 | 2.89 | 1.37 | 5.32$^{*}$ | 0.04 | 0.06 | 0.20 |
| RLW | leaf | 7.79$^{**}$ | 0.75 | 4.23$^{**}$ | 0.03 | 0.02 | 0.08 | 0.36 | 0.1 | 3.08 | 0.01 | 0.00 | 0.15 |
| RLL.RLW | leaf | 5.17$^{*}$ | 4.15$^{**}$ | 5.02$^{**}$ | 0.01 | 0.04 | 0.05 | 1.45 | 2.22 | 0.73 | 0.02 | 0.08 | 0.03 |
| SNLP | leaf | 27.17$^{***}$ | 1.51 | 8.08$^{***}$ | 0.10 | 0.03 | 0.12 | 16.53$^{**}$ | 2.32 | 4.85$^{*}$ | 0.20 | 0.09 | 0.18 |
| RNLP | leaf | 17.70$^{***}$ | 0.76 | 3.65$^{*}$ | 0.07 | 0.01 | 0.05 | 11.58$^{**}$ | 1.22 | 4.71$^{*}$ | 0.15 | 0.05 | 0.20 |

[1] F-values and corresponding significance levels in a generalized linear mixed-effect model are denoted. *P < 0.05, **P < 0.01, ***P < 0.001. Note only the last category passes the strict Bonferroni correction.
[2] Effect sizes (Eta2 = η2) of ecotype, region and their interaction estimated using the linear model

**Table S10** Morphological differentiation between foothill vs. alpine populations of *A. arenosa* quantified separately for each region using linear discriminant analysis (LDA). The analysis was based on in 16 traits collected in field samples and plants cultivated in common garden.

| | | Field data | | | | | | Common garden data | | | | |
|---|---|---|---|---|---|---|---|---|---|---|---|---|
| Trait | Category | All data | NT | VT | ZT | RD | FG | All data | NT | VT | ZT | FG |
| HP | stem | **0.74** | **-0.36** | **0.33** | **0.45** | **-0.61** | **0.78** | **-0.59** | **-0.39** | **-0.48** | **0.28** | **0.38** |
| HF | stem | **0.40** | 0.02 | **0.43** | **0.49** | **-0.42** | 0.16 | **-0.62** | **-0.39** | **-0.61** | **0.22** | **0.58** |
| PL | flower | **-0.28** | 0.07 | **-0.30** | -0.20 | **0.41** | -0.10 | **0.44** | **0.66** | **0.47** | -0.18 | -0.07 |
| PW | flower | **-0.40** | 0.13 | **0.39** | **0.48** | **0.35** | -0.18 | **0.56** | **0.79** | **0.47** | **-0.30** | -0.16 |
| PL.PW | flower | **0.26** | -0.10 | **0.44** | **0.64** | 0.03 | 0.14 | **-0.32** | **-0.49** | -0.09 | **0.22** | 0.19 |
| SL | flower | **-0.36** | 0.12 | -0.20 | **-0.30** | 0.16 | -0.12 | 0.23 | 0.16 | **0.37** | -0.11 | -0.01 |
| SW | flower | **-0.52** | 0.20 | **-0.60** | **-0.70** | 0.21 | **-0.33** | 0.12 | -0.07 | 0.25 | -0.14 | 0.05 |
| SL.SW | flower | **0.27** | -0.13 | **-0.60** | **-0.60** | -0.11 | **0.27** | 0.07 | 0.20 | 0.01 | 0.07 | -0.08 |
| SLL | leaf | 0.23 | 0.01 | **-0.30** | **-0.30** | 0.01 | **0.43** | 0.01 | 0.02 | 0.21 | -0.14 | 0.25 |
| SLW | leaf | 0.18 | 0.02 | **-0.40** | -0.01 | 0.17 | **0.35** | -0.16 | **-0.28** | 0.09 | -0.09 | **0.38** |
| SLL.SLW | leaf | 0.06 | -0.04 | 0.01 | -0.10 | **-0.32** | -0.07 | 0.21 | **0.29** | 0.06 | -0.06 | -0.16 |
| RLL | leaf | 0.08 | 0.02 | -0.01 | -0.20 | -0.16 | **0.28** | 0.14 | -0.01 | 0.03 | **-0.50** | 0.09 |
| RLW | leaf | 0.16 | -0.05 | -0.10 | -0.20 | -0.03 | **0.34** | 0.05 | -0.20 | 0.05 | **-0.40** | 0.14 |
| RLL.RLW | leaf | -0.10 | 0.10 | -0.10 | 0.03 | -0.25 | -0.06 | 0.10 | 0.10 | -0.01 | -0.21 | -0.01 |
| SNLP | leaf | **0.27** | -0.01 | 0.05 | -0.10 | 0.06 | **0.40** | **-0.33** | -0.15 | -0.17 | 0.04 | **0.64** |
| RNLP | leaf | **0.25** | -0.04 | 0.08 | -0.01 | 0.04 | 0.22 | **-0.31** | -0.05 | -0.24 | 0.03 | **0.33** |

Loadings on the first (constrained) axis of linear discriminant analysis (LDA) between foothill and alpine individuals calculated for the complete dataset ('all') and each region separately based on all 16 scored traits. The traits with moderate to high contribution to the discrimination (> 0.25 absolute value of the loading coefficient) are highlighted in bold.

**Supplementary data file 1** Observed allele frequency spectra, (**.dsfs* files), and example parameter files (**.est* and **.tpl* files) used for coalescent simulations in fastsimcoal

**Supplementary data file 2** Vegetation samples and environmental parameters sampled at original sites of *A. arenosa* populations

**Supplementary data file 3** Morphological traits of samples from field (sheet 1) and common garden (sheet 2).
